# BOLD signatures of sleep

**DOI:** 10.1101/531186

**Authors:** Chen Song, Melanie Boly, Enzo Tagliazucchi, Helmut Laufs, Giulio Tononi

## Abstract

Sleep can be distinguished from wake by changes in brain electrical activity, typically assessed using electroencephalography (EEG). The hallmark of non-rapid-eye-movement sleep are two major EEG events: slow waves and spindles. Here we sought to identify possible signatures of sleep in brain hemodynamic activity, using simultaneous fMRI-EEG. We found that, during the transition from wake to sleep, blood-oxygen-level-dependent (BOLD) activity evolved from a mixed-frequency pattern to one dominated by two distinct oscillations: a low-frequency (~0.05Hz) oscillation prominent in light sleep and a high-frequency (~0.17Hz) oscillation in deep sleep. The two BOLD oscillations correlated with the occurrences of spindles and slow waves, respectively. They were detectable across the whole brain, cortically and subcortically, but had different regional distributions and opposite onset patterns. These spontaneous BOLD oscillations provide fMRI signatures of basic sleep processes, which may be employed to study human sleep at spatial resolution and brain coverage not achievable using EEG.

**HIGHLIGHTS:** spontaneous BOLD oscillations differentiate sleep from wake

low-frequency BOLD oscillation tracks sleep spindles

high-frequency BOLD oscillation tracks sleep slow waves

BOLD oscillations provide fMRI signatures of key sleep processes

## INTRODUCTION

Brain activity during sleep is typically assessed using electroencephalography (EEG). The hallmark of non-rapid-eye-movement (NREM) sleep, which represents four fifths of human sleep, is the shift from high-frequency, low-amplitude EEG activity to low-frequency, high-amplitude EEG dominated by spindles and slow waves (Figure 1A). Spindles are characterized by waxing and waning rhythms in 11~16 Hz frequency range (herein referred to as sigma activity), which last for 1 to 3 seconds and occur around twice per minute (Zeitlhofer et al., 1997, Purcell et al., 2017). Slow waves are characterized by sharp negative deflections in 0.5~4 Hz frequency range (herein referred to as delta activity), which occur around seven times per minute (Mensen et al., 2016). Based on the prominence of spindles and slow waves, respectively, NREM sleep can be subdivided into a lighter N2 stage and a deeper N3 stage.

**Figure 1.**
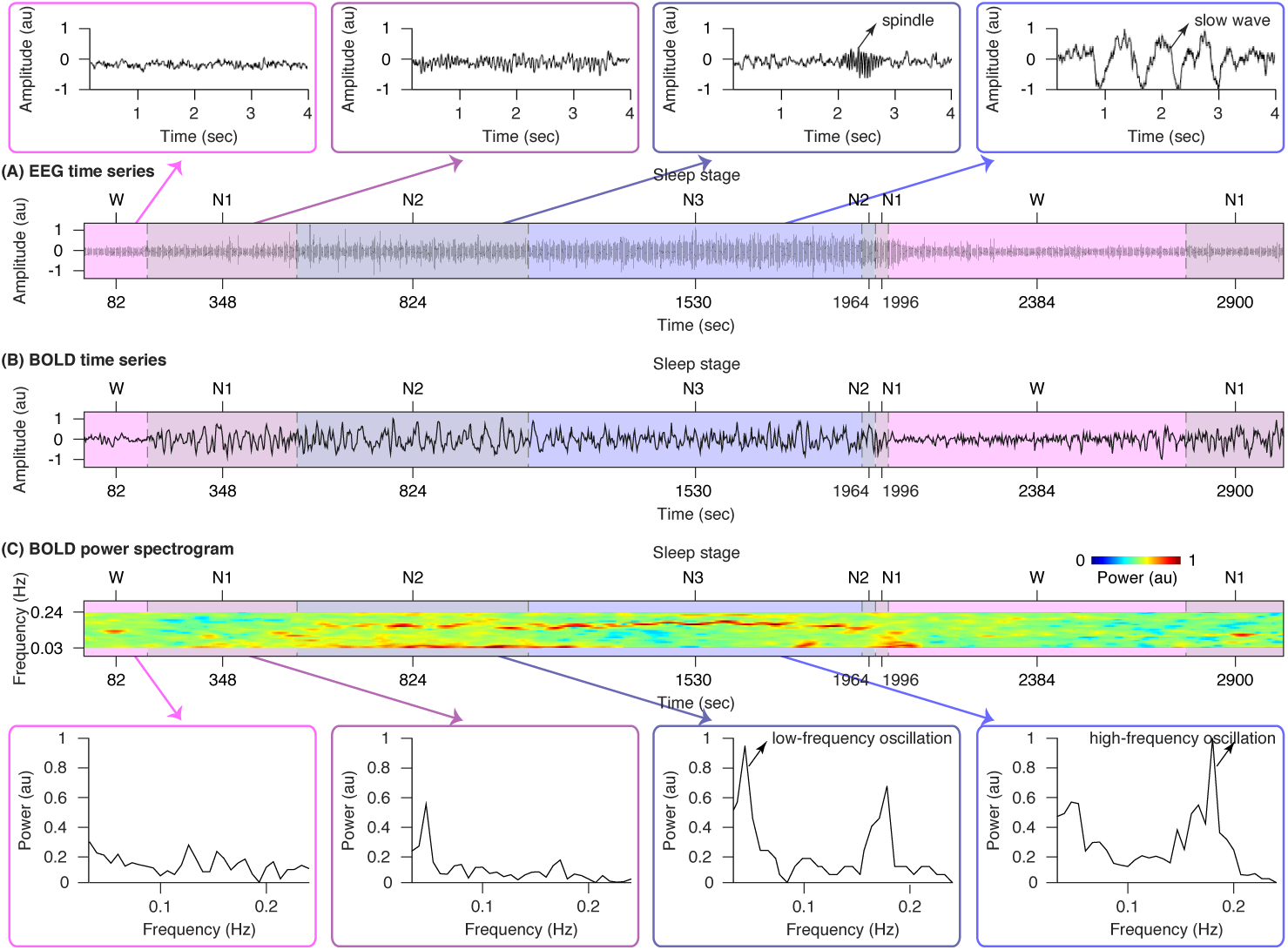
EEG and fMRI signatures of NREM sleep. (A) Representative EEG time series from simultaneous fMRI-EEG recording of sleep were plotted, illustrating the shift from high-frequency, low-amplitude wake EEG to low-frequency, high-amplitude sleep EEG dominated by spindles and slow waves. (B~C) Representative BOLD time series, power spectrogram and power spectrum from simultaneous fMRI-EEG recording of sleep were also plotted, illustrating the changes in BOLD frequency content from wake to sleep. During wakefulness, mixed-frequency, low-amplitude BOLD activity was observed. By contrast, during N1 then N2 sleep, BOLD activity evolved into a high-amplitude background with prominent low-frequency oscillation. Low-frequency BOLD oscillation attenuated when reaching N3 sleep, where most brain regions started to display spontaneous oscillations of higher frequency. The figure shows the data of participant S25.

Slow waves and spindles are being actively investigated because they are thought to mediate many of the restorative benefits of sleep at both a cellular and a systems level, including memory consolidation and integration (Rasch and Born, 2013, Tononi and Cirelli, 2014). Intracranial recordings have established that slow waves are associated with the near-synchronous transition in large populations of neurons between two distinct states, an up state of membrane depolarization and intense firing, and a down state of membrane hyperpolarization and silence (Steriade et al., 2001, Chauvette et al., 2011). Slow waves are generated primarily in the cortex (although the thalamus is required for their full expression), and affect virtually all cortical neurons, as well as neurons in several subcortical structures (Crunelli et al., 2015). By contrast, spindles are associated with cycles of hyperpolarization and depolarization triggered by interactions between reticular thalamic nucleus and specific thalamic nuclei, and further amplified by thalamo-cortico-thalamic circuits (Steriade et al., 1987). In natural sleep, slow waves may occur at different times in different cortical regions (Nir et al., 2011) and they can travel at a speed of a few meters per second over the cortical surface (Massimini et al., 2004, Murphy et al., 2009). Similarly, spindles exhibit regional specificity and show evidence of traveling (Andrillon et al., 2011, Piantoni et al., 2017, Hagler et al., 2018). Spindles and higher frequency EEG activities are often nested by underlying slow waves, and slow waves in turn can be grouped by infra-slow fluctuations (Steriade, 2000).

Although EEG has good temporal resolution, its spatial resolution is limited, strongly affected by volume conduction, and largely insensitive to neural activity in cortical and subcortical regions far away from the scalp. Here we sought to complement EEG recordings of human sleep by employing functional magnetic resonance imaging (fMRI) to investigate, with fuller brain coverage and greater spatial resolution, possible signatures of sleep in blood-oxygen-level-dependent (BOLD) activity. Spindles and slow waves are grouped by infra-slow fluctuations within the frequency range of hemodynamic responses (Chen and Glover, 2015, Lewis et al., 2016, Trapp et al., 2018). Therefore, while previous fMRI studies of sleep have mostly examined changes in BOLD amplitude (Schabus et al., 2007, Dang-Vu et al., 2008, Fukunaga et al., 2008, Picchioni et al., 2011, Schwalm et al., 2017) or functional connectivity (Laufs et al., 2007, Horovitz, et al., 2009, Larson-Prior et al., 2009, Boly et al., 2012, Duyn, 2012, Tagliazucchi et al., 2012, Liu et al., 2014, Picchioni et al., 2014), we investigated if the frequency content of BOLD activity shows systematic changes from wake to sleep, using simultaneous fMRI and polysomnographic EEG recordings.

We found that, during the transition from wake to sleep, BOLD activity evolved from a mixed-frequency pattern to one dominated by two distinct oscillations: a low-frequency (~0.05 Hz) oscillation that was prominent in light sleep and tracked the occurrence of spindles, and a high-frequency (~0.17 Hz) oscillation that was prominent in deep sleep and tracked the occurrence of slow waves. The low-frequency BOLD oscillation was strongest in sensory cortices and weaker in prefrontal cortex and subcortical regions, and it propagated from sensory to prefrontal cortices at the onset of sleep. By contrast, the high-frequency BOLD oscillation was strongest in prefrontal cortex and subcortical regions and weaker in sensory cortices, and it propagated from prefrontal to sensory cortices at sleep onset. The two spontaneous BOLD oscillations reported here provide fMRI signatures of basic sleep processes. They may be employed to study the occurrence, modulation, or function of sleep with higher spatial resolution than EEG and with the ability to probe deep brain structures.

## RESULTS

Simultaneous fMRI and polysomnographic EEG recordings were acquired from 58 non-sleep-deprived participants and constituted part of a larger dataset reported previously in (Tagliazucchi and Laufs, 2014). Recordings started at around 20:00 and lasted for 50 minutes. 36 out of 58 participants displaying sustained epochs of N2 and N3 sleep were included in subsequent analyses. The average time they spent in sleep was 33.27 minutes, and in N2 or N3 sleep was 22.80 minutes. The sleep architecture of all participants is summarized in Table S1.

### Changes in BOLD frequency content from wake to sleep

Representative EEG and BOLD time series from simultaneous fMRI-EEG recordings are displayed in Figure 1. Visual inspection of EEG time series suggested a shift from high-frequency, low-amplitude background during wake to low-frequency, high-amplitude background dominated by spindles and slow waves during sleep (Figure 1A). Visual inspection of BOLD time series suggested similar, frequency-specific changes starting at the transition from wake to sleep (Figure 1B). During wakefulness, mixed-frequency, low-amplitude BOLD activity was observed. By contrast, during N1 then N2 sleep, BOLD activity evolved into a high-amplitude background with prominent low-frequency oscillation. Low-frequency BOLD oscillation attenuated when reaching N3 sleep, where most brain regions started to display spontaneous oscillations of higher frequency.

To quantitatively assess the progressive changes in BOLD frequency content from wake to sleep, Fast Fourier Transform (FFT) analysis was applied to BOLD time series in sliding windows of 104 seconds (50 volumes) and step sizes of 2.08 seconds (1 volume, see Methods for details). The BOLD power spectrogram (Figure 1C) derived from FFT analysis confirmed our visual detection of a low-frequency BOLD oscillation emerging during N1 and N2 sleep, which progressively attenuated when transitioning to N3 sleep. It also revealed that the high-frequency BOLD oscillation visually detected in N3 sleep was already present, although at a lower power, during N2 sleep.

Based on the BOLD power spectrogram, we calculated the BOLD power spectrum for wake, N1, N2 and N3 sleep (Figure 1C). The BOLD power spectrum during wakefulness was relatively even, confirming visual impression of a mixed-frequency pattern. By contrast, the BOLD power spectrum during NREM sleep displayed two distinct and relatively sharp peaks: a first peak in lower-frequency range with a mean value of 0.05 Hz across participants (Table S2), and a second peak in higher-frequency range with a mean value of 0.17 Hz across participants (Table S3). The low-frequency peak appeared during the transition from wake to N1 sleep, became maximal in N2 sleep, and markedly attenuated in N3 sleep. By contrast, the high-frequency peak appeared in N2 sleep, and became maximal during N3 sleep. Consistent changes in BOLD frequency content from wake to sleep were observed across the whole brain, with the emergence of two distinct peaks in the power spectrum during sleep (Figure 2).

**Figure 2.**
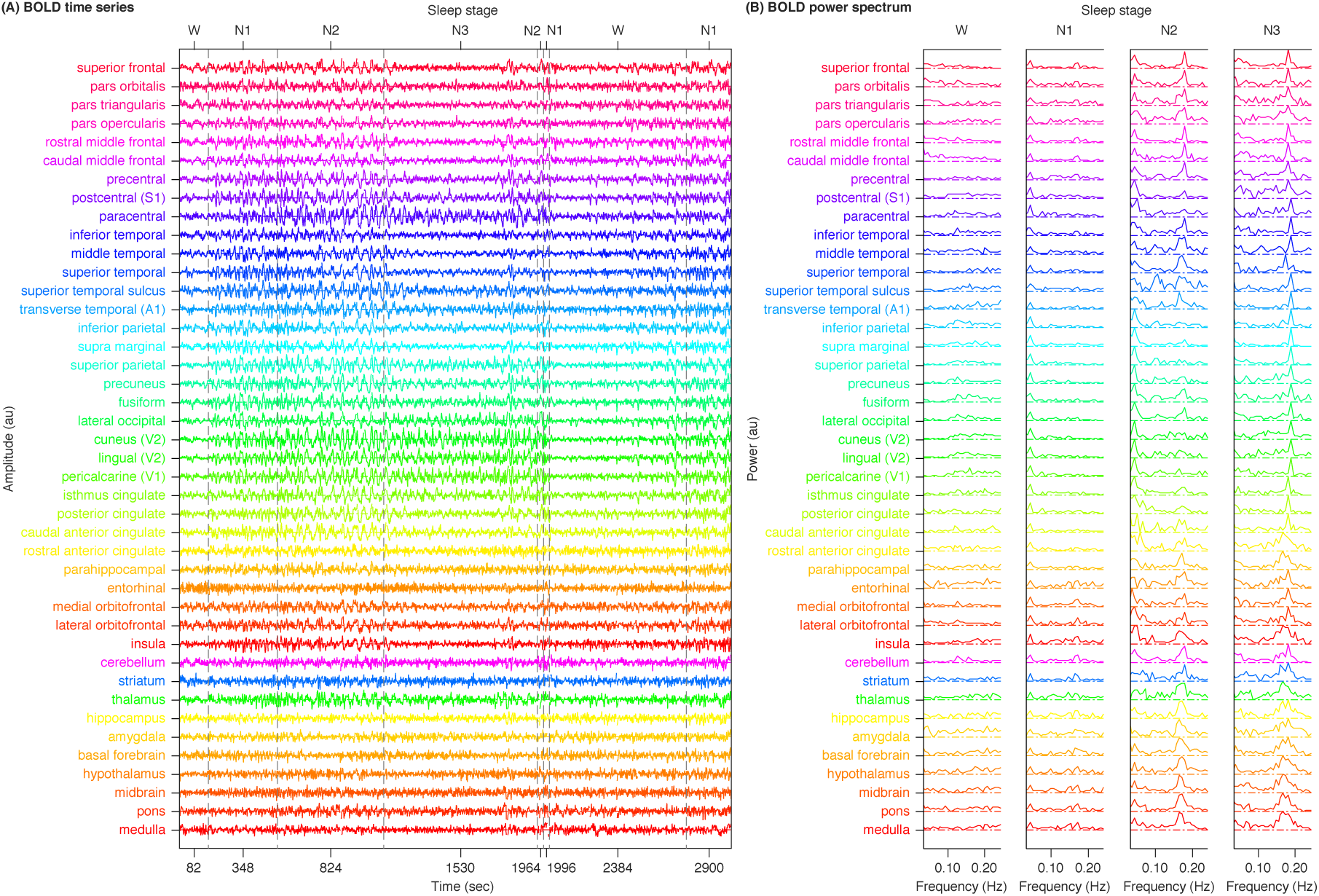
Whole-brain changes in BOLD frequency content from wake to sleep. Representative ROI-level BOLD time series and power spectrum (each color reflecting one ROI, see Figure 3A for color scales) were plotted, illustrating the consistency across different brain regions in the changes in BOLD frequency content from wake to NREM sleep. The figure shows the data of participant S25.

### Power topography of BOLD oscillations during sleep

To investigate the distribution of BOLD oscillations across the brain, we parcellated the cortex into 32 regions (Desikan et al., 2006) and the subcortex into 10 regions: cerebellum, striatum, thalamus, medulla, pons, midbrain, hypothalamus, basal forebrain, amygdala and hippocampus (Fischl et al., 2002, Zaborszky et al., 2008, Baroncini et al., 2012, Iglesias et al., 2016). In addition to this coarse atlas of 42 regions of interest (Figure 3A), we used a fine atlas of 217 regions of interest (Figure 3A), where the brain was parcellated into 180 cortical regions (Glasser et al., 2016), 11 cerebellum lobules (Diedrichsen et al., 2009), 7 striatum divisions (Tziortzi et al., 2014), 12 thalamic subregions (Figure S1; Morel et al., 1997, Krauth et al., 2010), medulla, pons, midbrain, hypothalamus, basal forebrain, amygdala and hippocampus (Fischl et al., 2002, Zaborszky et al., 2008, Baroncini et al., 2012, Iglesias et al., 2016). The number of voxels in individual regions of interest (ROIs) are listed in Table S4.

**Figure 3.**
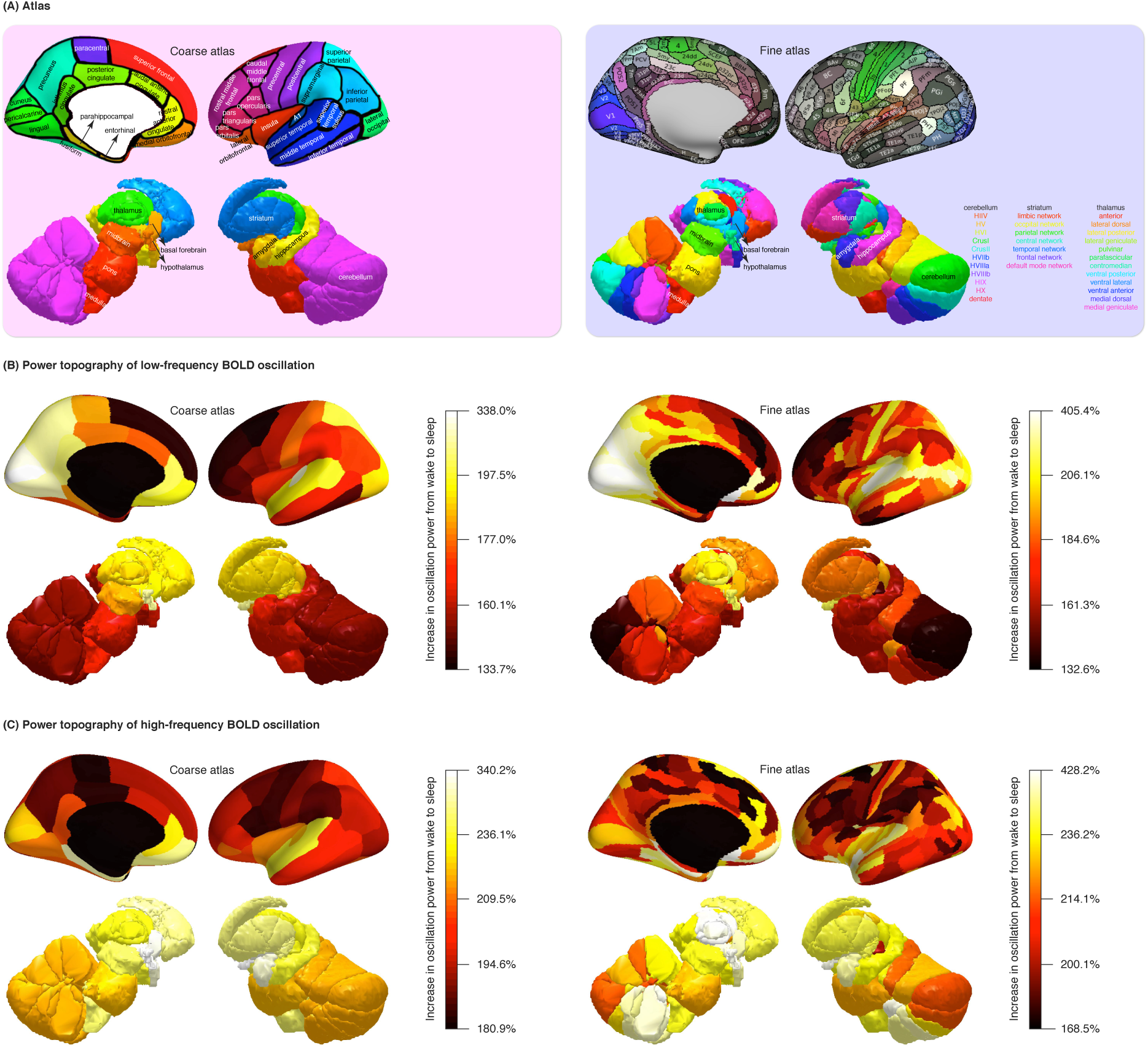
Power topography of BOLD oscillations. (A) Illustrated are the atlases used in our study, where the brain was parcellated into 42 coarse ROIs, including 32 cortical regions, cerebellum, striatum, thalamus, medulla, pons, midbrain, hypothalamus, basal forebrain, amygdala, hippocampus, or 217 fine ROIs, including 180 cortical regions, 11 cerebellum lobules, 7 striatum divisions, 12 thalamic subregions, medulla, pons, midbrain, hypothalamus, basal forebrain, amygdala, hippocampus. (B~C) Regional increases in BOLD oscillation power from wake to sleep were calculated. The power of low-frequency and high-frequency BOLD oscillations both showed a three-fold increase from wake to sleep. However, low-frequency BOLD oscillation (B) was generally strong in sensory and posteromedial cortices and weak in prefrontal and subcortical regions, whereas high-frequency BOLD oscillation (C) was strongest in prefrontal and subcortical regions and weaker in sensory and posteromedial cortices. Despite these differences, the subgenual area (an orbitofrontal region) had the strongest low-frequency BOLD oscillation as well as the strongest high-frequency BOLD oscillation among all cortical regions. The figure shows the average results across two hemispheres and participants S1 to S36.

We derived the power spectrogram of each ROI by applying FFT analysis on a voxel-level and calculating the average power spectrogram across all voxels within the ROI (see Methods for details). Representative ROI-level BOLD time series and power spectrum are displayed in Figure 2. On a participant-by-participant, ROI-by-ROI basis, we identified the spectral peaks in lower-frequency and higher-frequency ranges, traced the time courses of oscillation power, and calculated the increase in oscillation power from wake to sleep. The power of low-frequency and high-frequency BOLD oscillations both showed a two to three-fold increase from wake to sleep. Such an effect size is substantial, considering that the changes in BOLD amplitude evoked by stimuli or tasks during wakefulness are usually less than 10% of the baseline amplitude (Cohen et al., 2002, Sirotin et al., 2009). While they both increased several-fold from wake to sleep, we found that low-frequency (Figure 3B, Table S2) and high-frequency (Figure 3C, Table S3) BOLD oscillations had very different power topographies.

Within the cortex, low-frequency BOLD oscillation was strong in sensory and posteromedial regions (including visual areas V1, V2, V3, V3A, MT, V6, V6A, V7, V8, ProS, auditory areas A1, MBelt, LBelt, PBelt, RI, somatosensory areas 3a, 3b, precuneus and posterior cingulate area where the oscillation power increased by 304%, 266%, 230%, 244%, 248%, 348%, 273%, 237%, 225%, 257%, 259%, 293%, 282%, 269%, 230%, 207%, 209%, 250% and 261% from wake to sleep), but generally weak in prefrontal and parahippocampal regions (including Brodmann areas 8, 9, 10, 11, 32, 44, 45, 46, 47 and entorhinal cortex, where the oscillation power increased by 133%, 139%, 140%, 147%, 133%, 157%, 163%, 143%, 150% and 149%). By contrast, high-frequency BOLD oscillation was strong in prefrontal and parahippocampal regions (including Brodmann areas 10, 47 and entorhinal cortex, where the oscillation power increased by 285%, 266% and 303%), but weaker in sensory and posteromedial regions (including visual areas V2, V3, V3CD, MT, V4, V4t, V6, auditory areas A5, RI, somatosensory areas 2, 3a, 3b, precuneus and posterior cingulate area, where the oscillation power increased by 210%, 205%, 199%, 197%, 204%, 180%, 209%, 213%, 200%, 193%, 185%, 208%, 188% and 191%). Despite these differences in power topography, the subgenual area (an orbitofrontal region), had the strongest low-frequency BOLD oscillation (405% increase in oscillation power) as well as the strongest high-frequency BOLD oscillation (344% increase in oscillation power) among all cortical regions.

Within the subcortex, low-frequency BOLD oscillation was generally weaker than high-frequency BOLD oscillation. However, the two oscillations had similar power topography. Low-frequency BOLD oscillation was strongest in hypothalamus, basal forebrain and intralaminar, anterior, medial dorsal, lateral dorsal thalamus (where the oscillation power increased by 217%, 192%, 211%, 215%, 203% and 199%). Similarly, high-frequency BOLD oscillation was strongest in hypothalamus, basal forebrain and intralaminar, anterior, medial dorsal, lateral dorsal thalamus (where the oscillation power increased by 340%, 268%, 428%, 342%, 307% and 365%).

### BOLD oscillations track sleep spindles and slow waves

The two spontaneous BOLD oscillations observed during sleep, a low-frequency oscillation prominent in N2 sleep and a high-frequency oscillation in N3 sleep, seemed to mirror the well-known patterns of sleep EEG, with spindles (sigma activity) being prominent in N2 sleep and slow waves (delta activity) in N3 sleep (Figure 1). Moreover, the frequency of the two BOLD oscillations, 0.05 Hz and 0.17 Hz, was similar to the reported periodicity of spindles (about 2 times per minute, corresponding to 0.05 Hz, Zeitlhofer et al., 1997, Purcell et al., 2017) and slow waves (about 7 times per minutes, corresponding to 0.17 Hz, Mensen et al., 2016). Because of these similarities, we hypothesized that low-frequency and high-frequency BOLD oscillations might track spindle (sigma) and slow wave (delta) activities, respectively.

To test this hypothesis, we detected individual occurrences of spindles and slow waves using methods described in (Ferrarelli et al., 2007, Riedner et al., 2007). The time course of spindle or slow wave activity was calculated as the integral of their occurrence and duration in consecutive windows of 2.08 seconds (matching fMRI temporal resolution), and was correlated against the time course of low-frequency (Figure 4A) or high-frequency (Figure 5A) BOLD oscillation power, on a participant-by-participant, ROI-by-ROI basis. Since the intensity of spindle and slow wave activities can be approximated by the power of sigma (11~16 Hz) and delta (0.5~4 Hz) activities, respectively, we further derived sigma and delta power from EEG power spectrogram, and correlated the time course of sigma or delta activity against the time course of low-frequency (Figure 4B) or high-frequency (Figure 5B) BOLD oscillation power, on a participant-by-participant, ROI-by-ROI basis. The distributions of correlation coefficient across all participants and ROIs were plotted, where each value reflected the result from a single participant and a single ROI.

**Figure 4.**
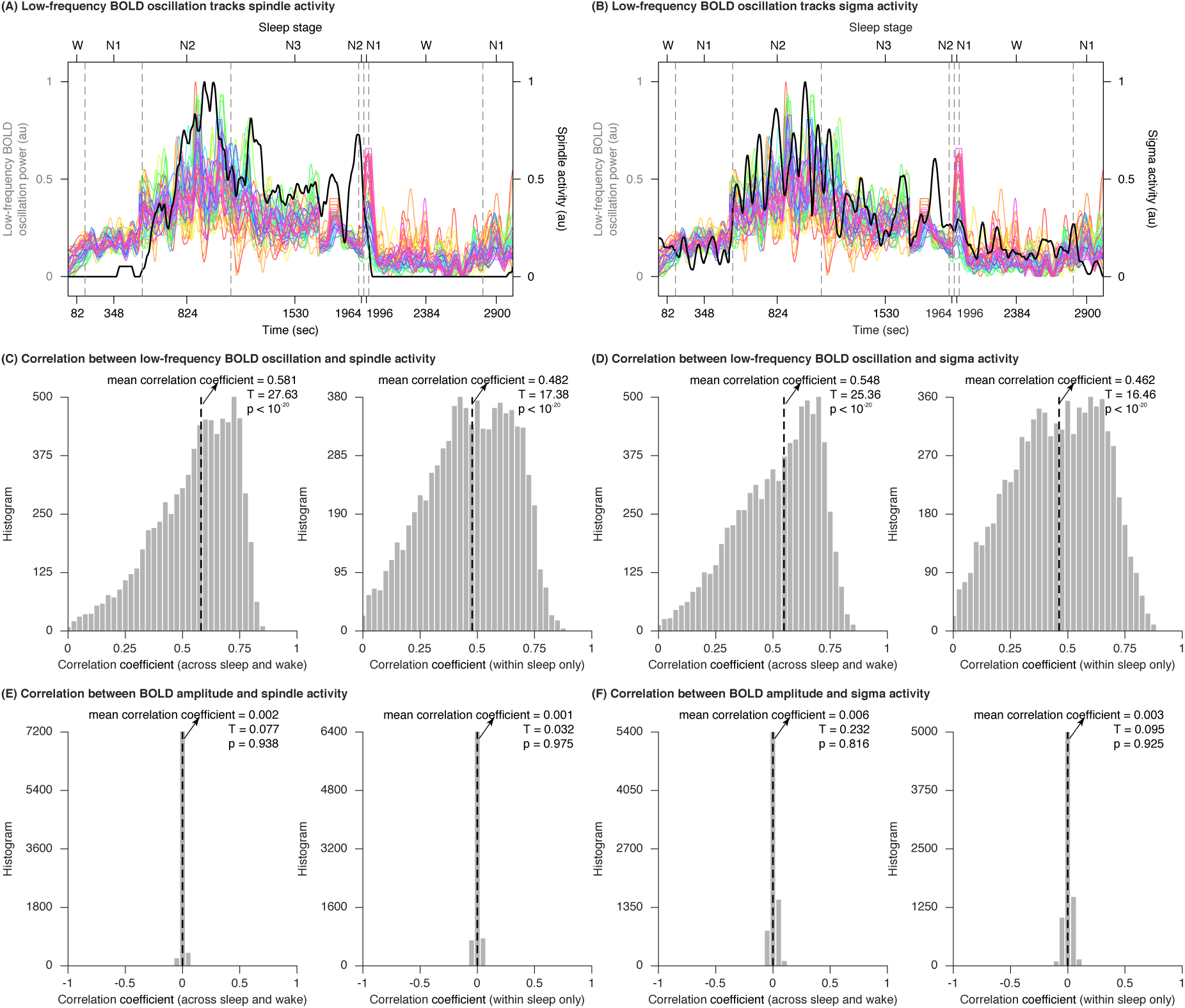
Low-frequency BOLD oscillation tracks sleep spindles. (A~B) Representative time courses of low-frequency BOLD oscillation power (colored lines, each color reflecting one ROI, see Figure 3A for color scales) and spindle or sigma activity (black lines) from simultaneous fMRI-EEG recording of sleep were plotted, illustrating the correlation between low-frequency BOLD oscillation and spindle or sigma activity. The figure shows the data of participant S25. (C~F) The time course of low-frequency BOLD oscillation power or BOLD amplitude was correlated against the time course of spindle or sigma activity, on a participant-by-participant, ROI-by-ROI basis, across sleep and wake (N = 1500 time points), or within sleep only (N = 1000 time points). The distributions of correlation coefficient across all participants and ROIs were plotted, where each value reflected the result from a single participant and a single ROI. At a data length of 1500 and 1000 time points, the threshold correlation coefficient for establishing statistical significance are 0.051 and 0.062 (before correction for multiple comparisons) or 0.136 and 0.166 (after Bonferroni correction for 374976 comparisons), respectively. The correlations between low-frequency BOLD oscillation power and spindle (C) or sigma (D) activity were much larger than the threshold correlation coefficient, and corresponded to T > 16.455, p < 10^-20^. By contrast, the correlations between BOLD amplitude and spindle (E) or sigma (F) activity were much smaller than the threshold correlation coefficient, and corresponded to T < 0.233, p > 0.815. The figure shows the average results across two hemispheres and participants S1 to S36.

**Figure 5.**
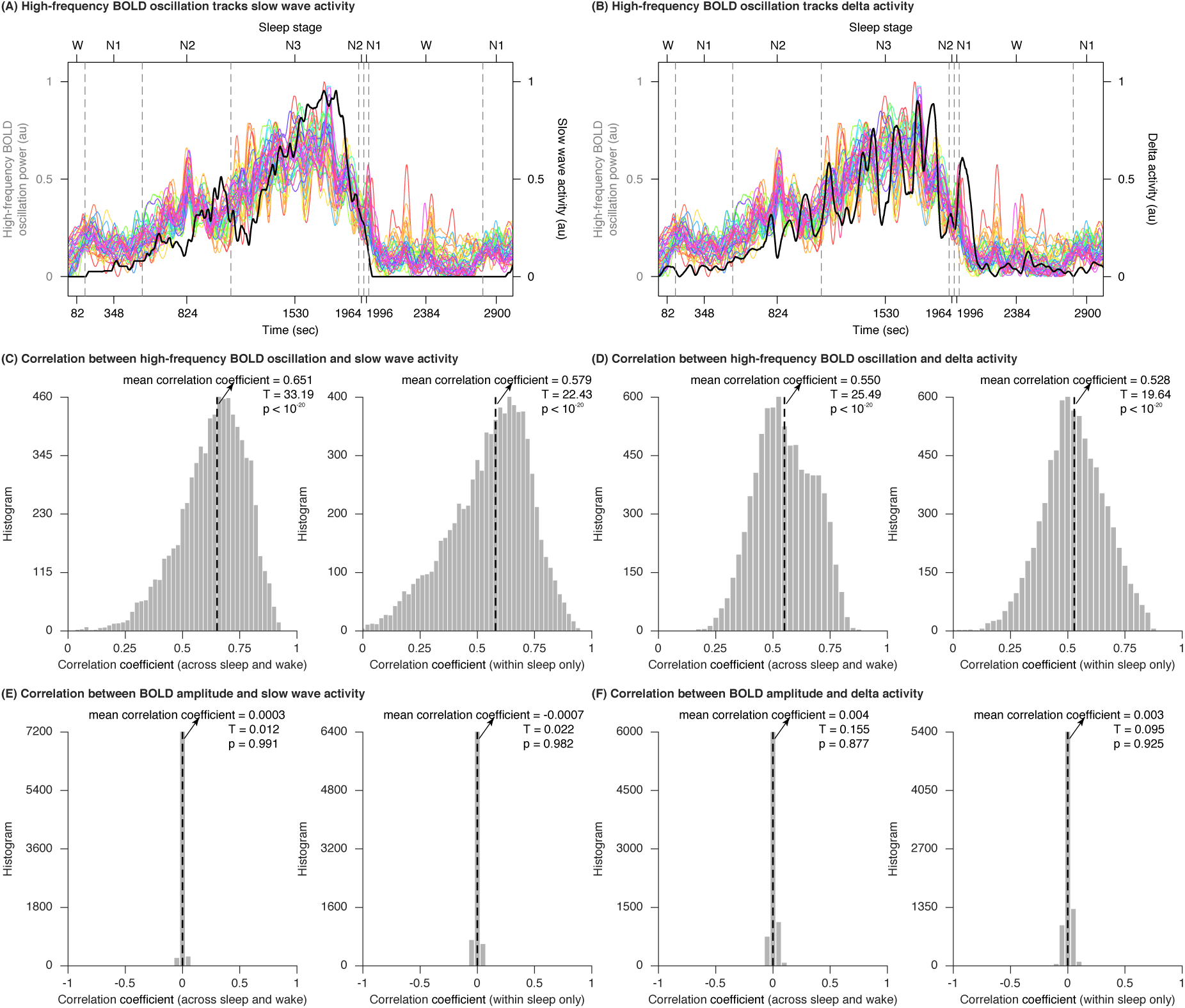
High-frequency BOLD oscillation tracks sleep slow waves. (A~B) Representative time courses of high-frequency BOLD oscillation power (colored lines, each color reflecting one ROI, see Figure 3A for color scales) and slow wave or delta activity (black lines) from simultaneous fMRI-EEG recording of sleep were plotted, illustrating the correlation between high-frequency BOLD oscillation and slow wave or delta activity. The figure shows the data of participant S25. (C~F) The time course of high-frequency BOLD oscillation power or BOLD amplitude was correlated against the time course of slow wave or delta activity, on a participant-by-participant, ROI-by-ROI basis, across sleep and wake (N = 1500 time points), or within sleep only (N = 1000 time points). The distributions of correlation coefficient across all participants and ROIs were plotted, where each value reflected the result from a single participant and a single ROI. At a data length of 1500 and 1000 time points, the threshold correlation coefficient for establishing statistical significance are 0.051 and 0.062 (before correction for multiple comparisons) or 0.136 and 0.166 (after Bonferroni correction for 374976 comparisons), respectively. The correlations between high-frequency BOLD oscillation power and slow wave (C) or delta (D) activity were much larger than the threshold correlation coefficient, and corresponded to T > 19.640, p < 10^-20^. By contrast, the correlations between BOLD amplitude and slow wave (E) or delta (F) activity were much smaller than the threshold correlation coefficient, and corresponded to T < 0.155, p > 0.876. The figure shows the average results across two hemispheres and participants S1 to S36.

The analysis revealed a strong correlation between low-frequency BOLD oscillation power and spindle activity (Figure 4C, within sleep only, r = 0.482, N = 1000 time points, T = 17.379, p < 10^-20^, across sleep and wake, r = 0.581, N = 1500 time points, T = 27.629, p < 10^-20^). It also revealed a strong correlation between high-frequency BOLD oscillation power and slow wave activity (Figure 5C, within sleep only, r = 0.579, N = 1000 time points, T = 22.434, p < 10^-20^, across sleep and wake, r = 0.651, N = 1500 time points, T = 33.193, p < 10^-20^). Moreover, low-frequency BOLD oscillation correlated more strongly with spindle than with slow wave activity (Figure S2A, within sleep only, ∆r = 0.302, N = 1000 time points, Z = 10.849, p < 10^-20^, across sleep and wake, ∆r = 0.313, N = 1500 time points, Z = 15.061, p < 10^-20^), whereas high-frequency BOLD oscillation correlated more strongly with slow wave than with spindle activity (Figure S2C, within sleep only, ∆r = 0.357, N = 1000 time points, Z = 13.742, p < 10^-20^, across sleep and wake, ∆r = 0.334, N = 1500 time points, Z = 17.362, p < 10^-20^).

Similar results were obtained for correlations between low-frequency BOLD oscillation power and sigma activity (Figure 4D, within sleep only, r = 0.462, N = 1000 time points, T = 16.457, p < 10^-20^, across sleep and wake, r = 0.548, N = 1500 time points, T = 25.356, p < 10^-20^), and between high-frequency BOLD oscillation power and delta activity (Figure 5D, within sleep only, r = 0.528, N = 1000 time points, T = 19.641, p < 10^-20^, across sleep and wake, r = 0.550, N = 1500 time points, T = 25.489, p < 10^-20^). Again, low-frequency BOLD oscillation correlated more strongly with sigma than with delta activity (Figure S2B, within sleep only, ∆r = 0.256, N = 1000 time points, Z = 9.184, p < 10^-20^, across sleep and wake, ∆r = 0.195, N = 1500 time points, Z = 9.543, p < 10^-20^), whereas high-frequency BOLD oscillation correlated more strongly with delta than with sigma activity (Figure S2D, within sleep only, ∆r = 0.368, N = 1000 time points, Z = 13.451, p < 10^-20^, across sleep and wake, ∆r = 0.272, N = 1500 time points, Z = 12.879, p < 10^-20^).

In contrast to the frequency content, the amplitude of BOLD activity did not show significant correlation with spindle activity (Figure 4E, within sleep only, r = 0.001, N = 1000 time points, T = 0.032, p = 0.975, across sleep and wake, r = 0.002, N = 1500 time points, T = 0.077, p = 0.938) or sigma activity (Figure 4F, within sleep only, r = 0.003, N = 1000 time points, T = 0.095, p = 0.925, across sleep and wake, r = 0.006, N = 1500 time points, T = 0.232, p = 0.816), or with slow wave activity (Figure 5E, within sleep only, r = −0.0007, N = 1000 time points, T = 0.022, p = 0.982, across sleep and wake, r = 0.0003, N = 1500 time points, T = 0.012, p = 0.991) or delta activity (Figure 5F, within sleep only, r = 0.003, N = 1000 time points, T = 0.095, p = 0.925, across sleep and wake, r = 0.004, N = 1500 time points, T = 0.155, p = 0.877). There was also no significant correlation between BOLD oscillation power and cardiac or respiratory activity (derived from respiration and pulse oximetry data), or cardiac or respiratory oscillation power (derived from FFT analysis to cardiac and respiratory time series) (Figure S3).

### Onset and offset of BOLD oscillations at wake-sleep transitions

Our findings suggest that low-frequency and high-frequency BOLD oscillations provide fMRI signatures of sleep spindles and slow waves, respectively. As such, the onset and offset of these BOLD oscillations may reflect which brain regions initiate the process of falling asleep and which brain regions are the first to wake up. To measure the onset and offset patterns of BOLD oscillations, we applied cross-correlation analysis to the time course of BOLD oscillation power and calculated the temporal lag between each pair of brain regions (see Methods for details, Mitra et al., 2014). The analysis was applied in sliding windows of 104 seconds and step sizes of 2.08 seconds, to capture the moment-to-moment changes in the lag structure. The average temporal lag that a brain region had with the rest of brain regions at the transition from wake to sleep (from 62.4 seconds before to 62.4 seconds after sleep onset) and at the transition from sleep to wake (from 62.4 seconds before to 62.4 seconds after sleep offset) indicated the relative timing of each region in the onset and offset of BOLD oscillations, respectively. The analysis revealed that the onset (Figure 6, Table S5) and offset (Figure 7, Table S6) of BOLD oscillations took place over a gradual course of about 10 seconds.

**Figure 6.**
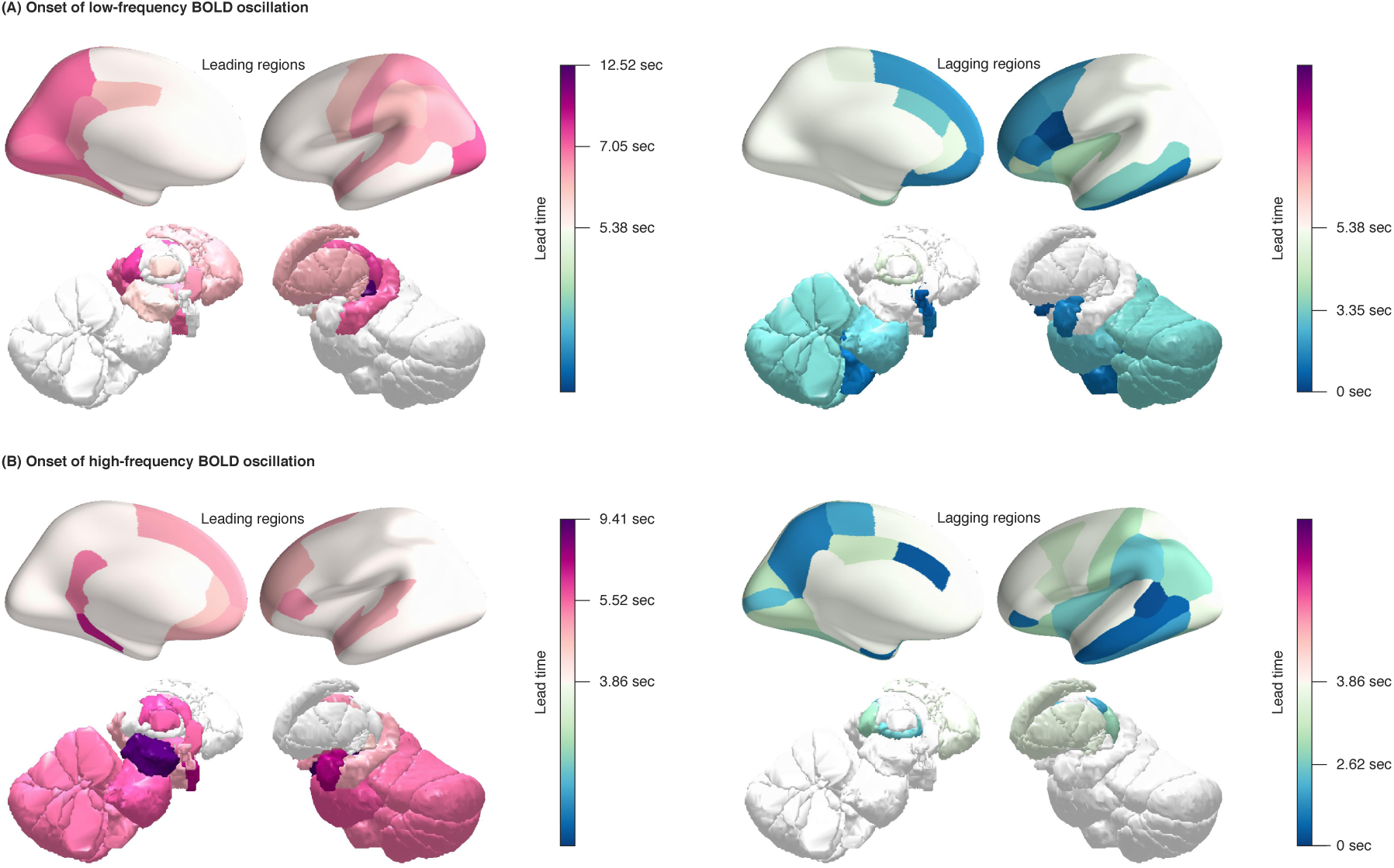
Onset of BOLD oscillations. The temporal lag between different brain regions in the onset of BOLD oscillations, at the transition from wake to sleep, was projected onto cortical and subcortical surfaces. Low-frequency (A) and high-frequency (B) BOLD oscillations had very different onset patterns: low-frequency BOLD oscillation first appeared in sensory thalamus, and propagated cortically from sensory to prefrontal regions; by contrast, high-frequency BOLD oscillation first appeared in midbrain, and propagated cortically from prefrontal to sensory regions. The figure shows the average results across two hemispheres and participants S1 to S36.

**Figure 7.**
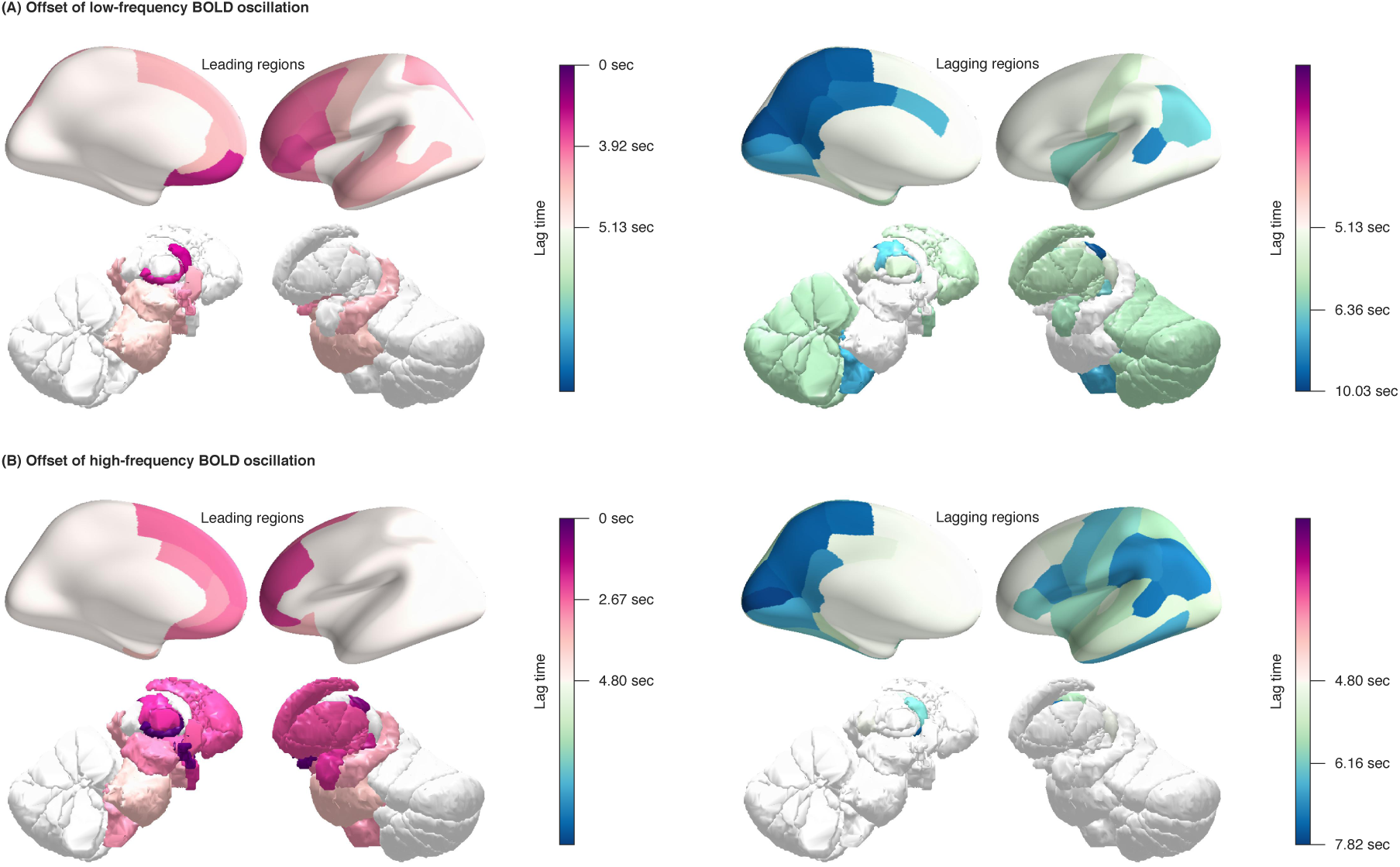
Offset of BOLD oscillations. The temporal lag between different brain regions in the offset of BOLD oscillations, at the transition from sleep to wake, was projected onto cortical and subcortical surfaces. Low-frequency (A) and high-frequency (B) BOLD oscillations had similar offset patterns: they both disappeared first from intralaminar thalamus, and their offsets propagated cortically from prefrontal to sensory regions. The figure shows the average results of two hemispheres across participants S1 to S36.

At the transition from wake to sleep, low-frequency BOLD oscillation first appeared in sensory thalamus (lateral geniculate and medial geniculate thalamus, where the oscillation led by 12.52 and 11.83 seconds), and then in other thalamic subregions (pulvinar, lateral posterior and ventral lateral thalamus, where the oscillation led by 8.88, 8.53 and 8.25 seconds). Soon after, the oscillation appeared in the cortex and propagated from sensory regions (V1, V2 and LOC, where the oscillation led by 7.87, 7.56 and 8.12 seconds) to prefrontal regions (superior frontal, middle frontal, orbitofrontal areas, pars triangularis and pars opercularis, where the oscillation led by 2.68, 2.74, 2.50, 2.32 and 0.99 seconds). By contrast, high-frequency BOLD oscillation first appeared in midbrain (9.41 seconds lead), followed by other subcortical regions (medial geniculate, lateral dorsal, medial dorsal, anterior, ventral lateral thalamus, amygdala, pons, medulla, basal forebrain and hypothalamus, where the oscillation led by 7.91, 6.44, 6.09, 6.19, 5.22, 7.63, 6.19, 5.55, 5.89 and 4.99 seconds). The oscillation then appeared in the cortex and propagated from prefrontal regions (isthmus cingulate, pars triangularis, superior frontal, middle frontal and orbitofrontal areas, where the oscillation led by 5.52, 5.17, 4.77, 4.29 and 4.52 seconds) to sensory regions (V1, inferior temporal, middle temporal areas, precuneus and entorhinal cortex, where the oscillation led by 2.34, 2.12, 1.01, 1.98 and 1.49 seconds).

Whereas low-frequency and high-frequency BOLD oscillations had nearly opposite onset patterns, they had similar offset patterns. At the transition from sleep to wake, low-frequency BOLD oscillation disappeared first from intralaminar thalamus (where the oscillation lagged by 0 seconds), and then from other thalamic subregions (anterior and medial geniculate thalamus, where the oscillation lagged by 1.62 and 3.11 seconds). Soon after, the oscillation disappeared in the cortex, along the direction from prefrontal regions (orbitofrontal, middle frontal areas, pars orbitalis, pars triangularis and pars opercularis, where the oscillation lagged by 2.22, 3.36, 2.81, 2.87 and 3.21 seconds) to sensory regions (V1, V2, superior temporal sulcus, posterior cingulate, isthmus cingulate, paracentral area and precuneus, where the oscillation lagged by 6.72, 7.74, 7.05, 7.44, 7.53, 7.54 and 7.65 seconds). Similarly, high-frequency BOLD oscillation disappeared first from intralaminar thalamus (0 seconds lag), followed by other thalamic subregions (lateral posterior, ventral posterior, lateral geniculate, medial dorsal and lateral dorsal thalamus, where the oscillation lagged by 0.83, 1.32, 1.99, 2.17 and 2.19 seconds). The oscillation then disappeared in the cortex, also along the direction from prefrontal regions (orbitofrontal, middle frontal, super frontal areas and pars orbitalis, where the oscillation lagged by 3.29, 2.54, 3.17 and 2.42 seconds) to sensory regions (inferior parietal area, superior temporal sulcus, paracentral area, precuneus, V2 and V1, where the oscillation lagged by 6.71, 6.86, 6.94, 7.04, 7.43 and 7.82 seconds).

## DISCUSSION

During the transition from wake to sleep, BOLD activity evolved from a mixed-frequency pattern to one dominated by two distinct oscillations: a low-frequency (~0.05 Hz) oscillation prominent during light sleep and a high-frequency (~0.17 Hz) oscillation prominent during deep sleep. The power of the two BOLD oscillations fluctuated within sleep and correlated with the fluctuations in spindle (sigma) and slow wave (delta) activities, respectively.

During sleep, low-frequency and high-frequency BOLD oscillations were detected across the whole brain, both cortically and subcortically, but with different power topographies: low-frequency BOLD oscillation was more prominent in sensory cortices, whereas high-frequency BOLD oscillation was strongest in subcortical regions and in prefrontal cortex. The two BOLD oscillations also had different onset patterns during the falling asleep process: low-frequency BOLD oscillation started in thalamus, then spread to sensory cortices, and finally to frontal cortex, whereas high-frequency BOLD oscillation started in midbrain, then spread to thalamus and frontal cortex, and finally to sensory cortices. The two oscillations had a similar offset pattern during the waking up process, disappearing first in thalamus, then in frontal cortex, and finally in sensory cortices.

The low-frequency and high-frequency BOLD oscillations reported here provide fMRI signatures of basic NREM sleep processes, which may be employed to study the mechanism and function of sleep at a higher spatial resolution and a fuller brain coverage than is achievable using EEG.

### A low-frequency BOLD oscillation is associated with sleep spindles

Our analysis revealed the emergence of a low-frequency BOLD oscillation during light sleep, which was correlated with the intensity of spindle activity. The peak frequency of this BOLD oscillation (~0.05 Hz) was similar to the reported periodicity of sleep spindles in the EEG (about 2 times per minute, corresponding to 0.05 Hz, Zeitlhofer et al., 1997, Purcell et al., 2017). While some distinguish between “fast” and “slow” spindles (Molle et al., 2011), this distinction remains controversial (de Genaro and Ferrara, 2003, D’Atri et al., 2017). Because BOLD signal displayed a single low-frequency peak in light sleep, we correlated its time course to broadband EEG spindle activity.

Previous studies have reported low-frequency BOLD oscillations during light sleep in human sensory cortices (Horovitz et al., 2008, Davis et al., 2016). The power of such BOLD oscillations was found to increase from birth to 1-2 years of age and correlate with the improvements in cognitive performance (Alcauter et al., 2015). Although previous studies did not examine the relations between such BOLD oscillations and sleep spindles, it is intriguing that sleep spindles emerge during the same age range (1-2 years old, Tanguay et al., 1975, Dan and Boyd, 2006). The co-emergence of low-frequency BOLD oscillation and EEG spindles during sleep in infancy may reflect the development of thalamo-cortical circuits key to both spindle generation (Steriade, 2000, D’Atri et al., 2017) and cognitive function (Alcauter et al., 2014).

Low-frequency BOLD oscillation was detected during sleep in all cortical and subcortical regions examined, consistent with the widespread detection of sleep spindles in intracranial recordings (Bazhenov et al., 2000, Andrillon et al., 2011, Nir et al., 2011). The power topography of low-frequency BOLD oscillation also mirrored that of spindle activity, with large-amplitude spindles reported intracranially in thalamus, sensory and orbitofrontal cortices (Bazhenov et al., 2000, Andrillon et al., 2011, Nir et al., 2011, Piantoni et al., 2017), and reduced occurrence of spindles in parahippocampal cortex (Andrillon et al., 2011). Moreover, the prominence of low-frequency BOLD oscillation in thalamus and sensory cortices is in line with previous EEG-fMRI studies of sleep spindles (Schabus et al., 2007, Caporro et al., 2012) and with a proposed role of spindles in sensory disconnection during sleep (Schabus et al., 2012).

### A high-frequency BOLD oscillation is associated with sleep slow waves

Spectral analysis of BOLD signal further revealed the emergence of a high-frequency BOLD oscillation during deep sleep, which was correlated with the intensity of slow wave activity. The peak frequency of this high-frequency BOLD oscillation (~0.17 Hz) was similar to the reported periodicity of EEG slow waves (about 7 times per minute, corresponding to 0.17 Hz, Mensen et al., 2016). A modulation of EEG slow waves can also occur at other slow EEG frequencies (Achermann and Borbely, 1997), but we did not find corresponding BOLD signal peaks during sleep.

Using direct current EEG recordings, a previous study demonstrated large-scale oscillations in the frequency range of 0.02 to 2 Hz during human sleep, which were synchronized with the occurrence of K complexes (Vanhatalo et al., 2004). Moreover, previous EEG-fMRI study reported correlations between slow BOLD fluctuation (<0.1 Hz) and EEG delta power during wake (Mantini et al., 2007), but changes between wake and sleep were not assessed. Finally, a study of propofol anesthesia found an increase in the mean frequency of BOLD activity compared to wake, with values approaching or surpassing 0.1 Hz (Guldenmund et al., 2016). While no correlation with EEG was performed in that study, propofol anesthesia can induce EEG slow waves similar to those of NREM sleep (Murphy et al., 2011).

High-frequency BOLD oscillation was observed during sleep in all cortical and subcortical regions examined. Similarly, intracranial recordings in humans and animals have demonstrated the ubiquitous occurrence of EEG slow waves in all cortical regions as well as in hippocampal formation and many subcortical regions (Contreras and Steriade, 1995, Steriade, 2006, Nir et al., 2011, Watson et al., 2016). The power topography of high-frequency BOLD oscillation also mirrored that of slow wave activity, and some of the regions showing the strongest high-frequency BOLD oscillation, including medial prefrontal cortex, parahippocampal cortex, and brainstem, were also highlighted in previous EEG-fMRI studies of sleep slow waves (Dang-Vu et al., 2008, Jahnke et al., 2012). The subgenual area, an orbitofrontal region, is notable for showing an especially strong sleep BOLD oscillation in both the low-frequency and the high-frequency range. Intriguingly, intracranial studies suggested that orbitofrontal cortex has among the highest amplitudes of sleep spindles and is involved in their propagation (Piantoni et al., 2017). Moreover, high-density EEG studies showed that orbitofrontal cortex is a hot spot for the origin and the propagation of sleep slow waves (Massimini et al., 2004, Murphy et al., 2009).

The power topography of high-frequency BOLD oscillation may already be interpreted with respect to several basic brain functions that can be influenced by the occurrence of slow waves. EEG slow waves reflect the synchronous down states across large populations of cells, which impairs neuronal responsiveness (Tononi and Massimini, 2008). Accordingly, the strong high-frequency BOLD oscillation in prefrontal cortex, which mirrors the large frontal slow wave activity, may explain the lack of responsiveness that is a hallmark of normal sleep (Boly et al., 2017). By contrast, the weak BOLD oscillation in several regions of the posterior cortex is consistent with the role of a posterior hot zone in supporting dream consciousness during two thirds of NREM sleep (Siclari et al., 2017). In turn, a strong BOLD oscillation in hippocampus and parahippocampal cortex may explain why most of these sleep dreams are quickly forgotten after awakening (Siclari et al., 2013). Based on the transient local decreases in high-frequency BOLD oscillation power, it may be possible to investigate which brain regions support specific contents of consciousness during sleep with much better spatial resolution than EEG. It will also be important to determine whether high-frequency BOLD oscillation may be present in primary sensory-motor areas during REM sleep (Chow et al., 2013), consistent with the recent demonstration of slow waves in these areas in both humans and animals (Funk et al., 2017, Baird et al., 2018) and potentially accounting for sensory-motor disconnection during an activated brain state.

### Onset and offset of low-frequency and high-frequency BOLD oscillations

The low-frequency and high-frequency BOLD oscillations observed during sleep differed not only in their power topography but also in their propagation patterns. At the transition from wake to sleep, thalamus was the first brain region to display low-frequency BOLD oscillation. This result is reminiscent of the intracranial finding that the inactivation of thalamus precedes that of the cortex during the falling asleep process (Magnin et al., 2010). In animals, reticular thalamic neurons have pacemaker properties that allow them to generate spindle-frequency oscillations through interactions with principal thalamic cells, especially within sensory relay thalamic nuclei (Sheroziya and Timofeev, 2014). The thin reticular thalamic nucleus could not be resolved in our fMRI data (Figure S1, Table S4), but the early onset of low-frequency BOLD oscillation in sensory thalamus (Figure 6, Table S5) is fully consistent with the cellular mechanism of spindle generation. Future studies using higher resolution EPI sequences are warranted to further investigate the differences in BOLD oscillatory behavior between different thalamic nuclei.

At the transition from wake to sleep, high-frequency BOLD oscillation was first initiated in the midbrain. Intriguingly, a unit recording study of midbrain reticular formation neurons in sleeping cats has found rhythmic fluctuations in neuronal firing rates with a modal period of 11 seconds (Oakson and Steriade, 1982, 1983), corresponding exactly to the BOLD oscillation frequency of ~0.17 Hz observed here during sleep. Such rhythmic fluctuations in midbrain firing rates were found to be phase-locked to increases in the amplitudes of cortical slow waves during sleep, but not during wake (Oakson and Steriade, 1983). Although we could not delineate midbrain reticular formation with our fMRI data, the frequency of the observed BOLD oscillation (~0.17 Hz) and its correlation with EEG slow wave activity are fully consistent with these unit recordings studies in cats. The fact that midbrain regions were the first to manifest high-frequency BOLD oscillation is also consistent with the central role of upper brainstem reticular formation in the regulation of normal arousal (Moruzzi and Magoun, 1949). An important issue is whether fluctuations in the firing of this midbrain region are responsible for modulating the excitability of the entire forebrain, thereby providing a common input that affects the likelihood of local and global slow waves (Bernardi et al., 2018). Another question is whether these midbrain firing fluctuations are generated endogenously and through which mechanisms.

Whereas low-frequency and high-frequency BOLD oscillations had very different onset patterns, they had similar offset patterns. At the transition from sleep to wake, intralaminar thalamus was the first brain region to show the offset of both low-frequency and high-frequency BOLD oscillations. During sleep, intralaminar thalamus also displayed a strong oscillation in both the low-frequency and the high-frequency range. In animals, electrical or chemical stimulation of intralaminar thalamic nuclei can reliably produce awakenings from sleep (Steriade and Demetrescu, 1960, 1961). During sleep, instead, the intralaminar thalamus is inhibited (Steriade et al., 1993). Human data also indicate that intralaminar thalamus is involved in the regulation of arousal as part of an anterior forebrain mesocircuit involving both subcortical and frontal regions (Saalmann, 2014, Schiff, 2010). In addition to intralaminar thalamus, other brain regions involved in arousal, such as hypothalamus and basal forebrain (Saper and Fuller, 2017, Scammel et al., 2017, Liu et al., 2018), were also among the first to show an offset of both low-frequency and high-frequency BOLD oscillations around the time of awakening.

### Potential mechanisms underlying BOLD oscillations in sleep

Overall, the present findings are highly consistent with previous evidence, obtained using several different approaches, about brain regions showing marked spindle activity or marked slow wave activity during sleep, respectively. They are also consistent with previous data about the onset and offset of spindles and slow waves. Our results suggest that the time courses of low-frequency and high-frequency BOLD oscillations were largely independent during sleep, with low-frequency BOLD oscillation correlating strongly with spindle activity and high-frequency BOLD oscillation with slow wave activity, but not the other way around. At the EEG level, an inverse temporal relationship between spindle activity and slow wave activity has often been reported, and their topography is also different (Werth et al., 1997, Tinguely et al., 2006, Purcell et al., 2017). On the other hand, sleep spindles are partly nested by sleep slow waves (Steriade, 2000), and both can be nested within lower frequency fluctuations. Thus, it is possible that the overall low-frequency and high-frequency fluctuations of excitability may have different, largely independent generators, but that the occurrence of individual slow wave up states may still increase the likelihood of spindle occurrence (Steriade, 2005).

The low-frequency and high-frequency BOLD oscillations were observed across the whole brain and provide fMRI signatures of sleep spindles and slow waves, respectively. However, it should be emphasized that the detection of local BOLD oscillation during sleep within a brain region cannot be expected to have a one-to-one correspondence with the occurrence of local EEG spindles or slow waves, but at best to reflect the grouping of spindles and slow waves by lower frequency fluctuations. In our study, both low-frequency and high-frequency BOLD oscillations were detected in subcortical regions, such as cerebellum and brainstem, where spindles and slow waves have not previously been recorded using either local EEG or unit recordings. Three main possibilities should be considered. First, it is possible that the presence of a BOLD oscillation is a signature of the occurrence of spindles or slow waves that had not been investigated or detected. This possibility is most likely for cortical regions whose sleep-related activity has not yet been documented, especially in the human brain. Slow fluctuations in BOLD signal can relate to local fluctuations in neuronal firing (Kim et al., 2004, Nir et al., 2007), and in isoflurane-anesthetized rats BOLD signal traced neuronal firing time-locked to EEG slow waves (Schwalm et al., 2017). Conceivably, the reduced energy needs during slow wave down states, related to reduced glial energetic demands for the repolarization cell membranes after action potentials (Wells et al., 2015), may be partly responsible for the correlation between the power of BOLD oscillation and the occurrence of sleep slow waves.

A second possibility is that low-frequency and high-frequency BOLD oscillations detected during sleep may reflect fluctuations in neuronal excitability that do not necessarily result in EEG spindles or slow waves. For example, diffuse subcortical projections from midbrain reticular formation may not only affect the likelihood of up and down states in the cortex, leading to changes in EEG slow wave activity, but do so also in brainstem or cerebellar regions where neurons may lack the mechanisms for producing up and down states.

As a third possibility, in regions lacking such mechanisms for spindle or slow wave generation, the low-frequency and high-frequency BOLD oscillations may reflect synaptic input synchronized with the occurrence of spindles or slow waves elsewhere in the brain. The cortical slow waves, which involve the synchronous down states across large populations of neurons, are bound to impose a strong input modulation onto many subcortical targets, directly or indirectly, starting with basal ganglia circuits and including brainstem and cerebellum. As a case in point, most neurons in the hippocampus lack the ability to produce down states (Isomura et al., 2006), but they show a strong modulation in local field potentials in the slow oscillation range originated in the cortex (Molle et al., 2006). Accordingly, animal experiments have shown that fluctuations in BOLD signal correlate better with local field potentials than with neuronal firing (Logothetis et al., 2001, Ureshi et al., 2004, Bartolo et al., 2011, Thompson et al., 2014, Murta et al., 2015), and regional increases in cerebral blood flow can occur even in the absence of neuronal spiking activity (Mathiesen et al., 2000). Thus, the detection of BOLD oscillations in regions lacking mechanisms for spindle or slow wave generation may reflect the modulation from other brain regions.

In our study, there was no significant correlation between BOLD oscillation power and cardiac or respiratory activity, suggesting that the present findings are unlikely to be explained by non-neuronal physiological effects. Nonetheless, BOLD signal results from a complex interplay between neuronal and vascular events (Friston et al., 2000, Obrig et al., 2000, Logothetis et al., 2001, Shmueli et al., 2007, Bianciardi et al., 2009, Webb et al., 2013, Yuan et al., 2013, Rayshubskiy et al., 2014, Chang et al, 2016, Tong et al., 2016), where oscillations in the signal could be affected by vasomotion (~0.1 Hz intrinsic fluctuations in arteriole diameter) or by autonomous nervous system activity (cardiac or respiratory activity). Notably, dynamic coupling between cardiac activity and spindle or slow wave activity has been reported during human sleep (Lechinger et al., 2015, Lin et al., 2016, Menson et al., 2016, de Zambotti et al., 2018), which predicts the post-sleep improvements in cognitive performances (Naji et al., 2018), consistent with the visceral influences on brain and behavior observed during wake (Critchley and Harrison, 2013, Park et al., 2014). Moreover, vasomotion is entrained by neuronal oscillations of similar frequency (Mateo et al., 2017). This entrainment is considered to be the basis of resting-state BOLD fMRI (He et al., 2018). It also underlies the low-frequency (~0.06 Hz) and high-frequency (~0.12 Hz) hemodynamic oscillations observed in anesthetized rats and the correlations of these hemodynamic oscillations with intracellular calcium oscillation (~0.07 Hz) (Du et al., 2014). As such, vasomotion and autonomous nervous system activity may provide the physiological links between neuronal and hemodynamic signals. Filtering out BOLD frequencies >0.1 Hz, as routinely applied in fMRI studies of resting-state, may discard neuronal signals of interest (Chen and Glover, 2015, Lewis et al., 2016, Trapp et al., 2018), especially when high-frequency BOLD oscillation is prominent as in deep sleep.

Limited by the temporal resolution of fMRI, we could not study how the propagation of BOLD oscillations relates to that of individual spindles or slow waves, which travel at a speed of a few meters per second (Massimini et al., 2004, Murphy et al., 2009, Andrillon et al., 2011, Piantoni et al., 2017, Hagler et al., 2018). Because sleep fMRI recordings only lasted an hour, we could also not assess properly whether the power of BOLD oscillations changes during the course of sleep, or if the parameters of BOLD oscillations exhibit trait-like, inter-individual variability. It is well known that sleep slow wave activity is homeostatically regulated, decreasing exponentially over the course of a night (Borbély, 1982, Riedner et al., 2007, Dijk, 2009), and that the overnight changes in slow wave activity correlate with the improvements in cognitive performances after sleep (Huber et al., 2004, 2006, Mascetti et al., 2013, Bellesi et al., 2014, Bernardi et al., 2015). Spindles, on the other hand, exhibit trait-like variability across individuals and correlate with individual differences in cognitive abilities (Bodizs et al., 2005, Schabus et al., 2006, Fogel et al., 2007, Geiger et al., 2011). Compared to EEG measures of spindles or slow waves, fMRI measures of BOLD oscillations have much higher spatial resolution and fuller brain coverage. The spatial topography of BOLD oscillations may allow a finer characterization of the links between sleep and cognition than is possible using EEG.

## ACKNOWLEDGEMENTS

The study was supported by Tiny Blue Dot Foundation grant (to GT), Wellcome Trust (209192/Z/17/Z to CS), H2020 MSCA COFUND (663830-CU119 to CS), Bundesministerium für Bildung und Forschung (grant 01 EV 0703 to ET and HL) and LOEWE Neuronale Koordination Forschungsschwerpunkt Frankfurt (NeFF to ET and HL). We thank Chiara Cirelli, Brady Riedner and Armand Mensen for their help and suggestions.

## METHODS

### Participants

Simultaneous fMRI and polysomnographic EEG recordings were acquired from 58 non-sleep-deprived participants and constituted part of a large dataset reported previously in (Tagliazucchi and Laufs, 2014). The recordings started at around 20:00 and lasted for 52 minutes, in a Siemens Trio 3T MRI scanner with an optimized polysomnographic setting. Participants were instructed to close their eyes, lie still and relaxed in the scanner. Written informed consent was obtained from all participants and the study was approved by Goethe University Ethics Committee. 36 out of 58 participants displaying sustained epochs of N2 and N3 sleep were included in subsequent analyses (Table S1).

### Data acquisition and preprocessing

Polysomnographic EEG data were acquired using a 30-channel BrainCap MR (sampling rate: 5 kHz; reference channel: FCz), and preprocessed in BrainVision Analyzer (www.brainproducts.com) for MRI and pulse artifact correction (average artifact subtraction method and ICA-based rejection of residual artifact-laden components). The preprocessed data were down-sampled to 128 Hz and re-referenced to the average of all channels. Other polysomnographic data included electrooculography (EOG), electromyography (EMG), electrocardiography (ECG), respiration and pulse oximetry. Sleep staging was performed according to the American Academy of Sleep Medicine (AASM) criteria.

Functional MRI data were acquired using a T2*-weighted EPI sequence (TR: 2.08 seconds; TE: 30 msec; field of view: 192 x 192 x 128 mm; resolution: 3 x 3 x 2 mm), and preprocessed in SPM (http://www.fil.ion.ucl.ac.uk/spm) using realignment, unwarping, and correction for head motion, cardiac activity and respiratory activity (RETROICOR method, Glover et al., 2000). The preprocessed data were high-pass filtered (cut-off frequency: 0.03 Hz) to remove slow signal drift.

Structural MRI data were acquired using a T1-weighted sequence (TR: 14.25 msec; TE: 5.99 msec; field of view: 224 x 256 x 176 mm; resolution: 1 mm isotropic), and preprocessed in Freesurfer (https://surfer.nmr.mgh.harvard.edu/) using non-uniform intensity correction, skull-stripping, and tissue segmentation into gray matter, white matter and cerebrospinal fluid. The white mater and gray matter segments were covered with triangular tessellations to reconstruct the three-dimensional white and pial surfaces.

### EEG spindle and slow wave activities

Individual occurrences of sleep spindles and slow waves were detected using methods described in (Ferrarelli et al., 2007, Riedner et al., 2007). For spindle detection, EEG time series were band-pass filtered at 11~16 Hz (Chebyshev Type II filter), and spindles with a duration of 0.3~3 seconds and an amplitude higher than eight standard deviations of the mean were detected. For slow wave detection, EEG time series were band-pass filtered at 0.5~4 Hz (Chebyshev Type II filter), and slow waves with a duration of 0.25~1.25 seconds and an amplitude higher than five standard deviations of the mean were detected.

The filter parameters were optimized to achieve minimal wave shape and amplitude distortion while allowing the least high-frequency contamination. The detection thresholds were based on Rechtschaffen and Kales criterion (Rechtschaffen and Kales, 1968) and on previous studies comparing different detection methods (Uchida et al., 1999, Geering et al., 1993, Warby et al., 2014). The results of automated detections were manually checked. Then, the time courses of spindle and slow wave activities were calculated as the integral of their occurrence and duration in consecutive windows of 2.08 seconds (matching fMRI temporal resolution).

### EEG sigma and delta activities

Since spindle and slow wave activities are reflected in the sleep EEG by sigma (11~16 Hz) and delta (0.5~4 Hz) activities, respectively, we further used the time courses of sigma and delta activities to approximate the time courses of spindle and slow wave activities. To this end, Fast Fourier Transform (FFT) analysis was applied to EEG time series in sliding windows of 104 seconds and step sizes of 2.08 seconds (matching fMRI temporal resolution), and the time courses of sigma and delta power were derived from EEG power spectrogram.

### MRI brain parcellation

Based on the morphology (sulcus, gyrus) of reconstructed cortical surfaces, the cortex was parcellated into 32 regions (Desikan et al., 2006), and based on probabilistic histological atlas, the subcortex was parcellated into 10 regions: cerebellum, striatum, thalamus, medulla, pons, midbrain, hypothalamus, basal forebrain, amygdala and hippocampus (Fischl et al., 2002, Zaborszky et al., 2008, Baroncini et al., 2012, Iglesias et al., 2016). In addition to this coarse atlas of 42 regions of interest (Figure 3A), we used a fine atlas of 217 regions of interest (Figure 3A), in which the brain was parcellated into 180 cortical regions (Glasser et al., 2016), 11 cerebellum lobules (Diedrichsen et al., 2009), 7 striatum divisions (Tziortzi et al., 2014), 12 thalamic subregions (Figure S1; Morel et al., 1997, Krauth et al., 2010), medulla, pons, midbrain, hypothalamus, basal forebrain, amygdala and hippocampus (Fischl et al., 2002, Zaborszky et al., 2008, Baroncini et al., 2012, Iglesias et al., 2016).

The fine parcellations of cerebellum, striatum and thalamus were performed through spatial transformation of the brain atlas (in MNI152 space) to individual participants’ structural MRI data (in subject-native space) using SPM. The rest of the parcellations were performed in subject-native space using Freesurfer. The results of brain parcellation, performed on the higher-resolution structural MRI data, were co-registered and re-sliced to the lower-resolution functional MRI data.

### MRI spectral analysis

Fast Fourier Transform (FFT) analysis was applied to fMRI BOLD time series in sliding windows of 104 seconds (50 volumes) and step sizes of 2.08 seconds (1 volume) on a voxel-level (to account for the potential differences across voxels in the phase of BOLD time series). The power spectrogram of each region of interest (ROI) was calculated as the average power spectrogram across all voxels within this ROI, normalized by their total power (Baria et al., 2011).

The normalization procedure was conceptually similar to the one usually taken in deriving ROI-level BOLD time series from voxel-level BOLD time series; it was essential because BOLD signal is a qualitative and relative measure rather than a quantitative and absolute measure (Griffeth et al., 2011). As such, the abolute value of BOLD oscillation power is not biologically interpretable or meaningful; only the relative values, such as the increase in BOLD oscillation power from wake to sleep, or the fluctuations in BOLD oscillation power within sleep, are biologically interpretable. These relative measures were used in subsequent analysis.

On a participant-by-participant, ROI-by-ROI basis, we identified the spectral peaks in lower-frequency and higher-frequency ranges, and traced the time courses of BOLD oscillation power. Based on the time courses, we calculated the increase in BOLD oscillation power from wake to sleep and the statistical difference (T value) in BOLD oscillation power between wake and sleep (Figure 3, Table S2, Table S3). The time courses of BOLD oscillation power were also used to calculate the correlation between BOLD oscillations and spindle, sigma, slow wave, delta, cardiac or respiratory activity (Figure 4, Figure 5, Figure S2, Figure S3). Moreover, cross-correlation analysis was applied to the time courses of BOLD oscillation power, to estimate the temporal lag between different brain regions in the onset or offset of BOLD oscillations (Figure 6, Figure 7, Table S5, Table S6). The details of these analyses are described below.

### Correlation between BOLD oscillations and EEG spindle and slow wave activities

The time course of low-frequency BOLD oscillation power, high-frequency BOLD oscillation power, or BOLD amplitude, was correlated against the time course of spindle, sigma, slow wave or delta activity, on a participant-by-participant, ROI-by-ROI basis, across sleep and wake (N = 1500 time points), or within sleep only (N = 1000 time points). Moreover, the time course of BOLD oscillation power was correlated against the time course of cardiac and respiratory activities (derived from respiration and pulse oximetry data) or the time course of cardiac and respiratory oscillation power (derived from FFT analysis to cardiac and respiratory time series). The distributions of correlation coefficient (Figure 4, Figure 5, Figure S3), or the distributions of difference in correlation coefficient (Figure S2), across all participants and ROIs were plotted, where each value reflected the result from a single participant and a single ROI.

From the correlation coefficient (r), the statistical significance (T value, p value) can be calculated as: T=sqrt(N-2)*r/sqrt(1-r^2^), p=2*tcdf(T,N-2), where N is the data length, N-2 is the degree of freedom, and tcdf is the T cumulative distribution function. At a data length of 1500 and 1000 time points, the threshold correlation coefficient for establishing statistical significance are 0.051 and 0.062, respectively, before correction for multiple comparisons, or 0.136 and 0.166, respectively, after Bonferroni correction for 374976 comparisons, composed of 16 variables (spindle, slow wave, sigma, delta, 6 cardiac, 6 respiratory) * 3 variables (low-frequency BOLD oscillation power, high-frequency BOLD oscillation power, BOLD amplitude) * 217 ROIs * 36 participants.

The statistical significance (Z value, p value) for the difference in correlation coefficient (∆r) can be calculated as: Z_1_=0.5*(log(1+r_1_)-log(1-r_1_)), Z_2_=0.5*(log(1+r_2_)-log(1-r_2_)), Z=(Z_1_-Z_2_)/sqrt(1/(N-3)), p=2*normcdf(Z,0,1), where r_1_ and r_2_ are the two correlation coefficient values, Z_1_ and Z_2_ are their corresponding Z values, N is the data length, N-3 is the degree of freedom, and normcdf is the normal cumulative distribution function.

### Onset and offset of BOLD oscillations

Cross-correlation analysis was applied to estimate the temporal lag between different brain regions in the onset or offset of BOLD oscillations, on a participant-by-participant basis. For each pair of brain regions, cross-correlation between the time course of their BOLD oscillation power was calculated and was interpolated with parabolic function to quantify the temporal lag at which the correlation coefficient was maximal (Mitra et al., 2014). The interpolation procedure allowed the temporal lag to be estimated at a resolution finer than fMRI temporal resolution. Based on the pair-wise calculation, a lag matrix was derived, in which individual columns reflected the temporal lag that individual brain regions had with the rest brain regions.

The cross-correlation analysis was applied in sliding windows of 104 seconds and step sizes of 2.08 seconds, to capture the moment-to-moment changes in the lag structure. The average temporal lag that a brain region had with the rest brain regions, at the transition from wake to sleep (from 62.4 seconds before to 62.4 seconds after sleep onset), and the transition from sleep to wake (from 62.4 seconds before to 62.4 seconds after sleep offset), quantified the relative position of this region in the onset (Figure 6, Table S5) and offset (Figure 7, Table S6) of BOLD oscillations, respectively.

