## Supplemental Figures S1-S3 & Tables S1-S6 for "BOLD signatures of sleep"

### SUPPLEMENTAL MATERIALS

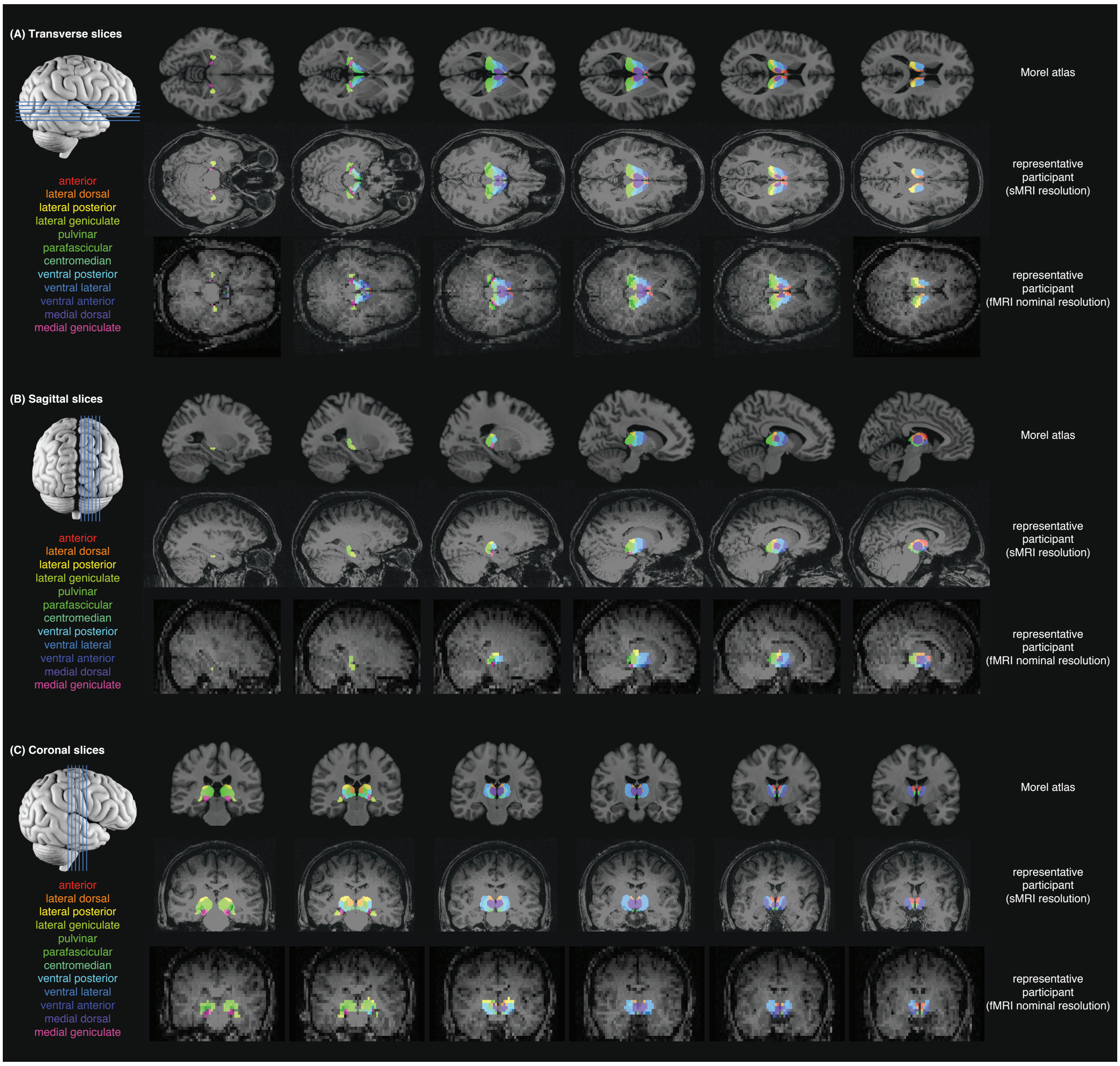


**Figure S1. Thalamus parcellation.** Based on Morel atlas (Morel et al., 1997, Krauth et al., 2010), the thalamus was parcellated into anterior, lateral dorsal, lateral posterior, lateral geniculate, pulvinar, parafascicular, centromedian, ventral posterior, ventral lateral, ventral anterior, medial dorsal, and medial geniculate subregions. The parcellation was performed through spatial transformation of Morel atlas (in MNI152 space) to individual participants' structural MRI data (in subject-native space). The results of thalamus parcellation, performed on the higher-resolution structural MRI data, were then co-registered and re-sliced to the lower-resolution functional MRI data. Illustrated are the Morel atlas in MNI152 space and the parcellation results of a representative participant.

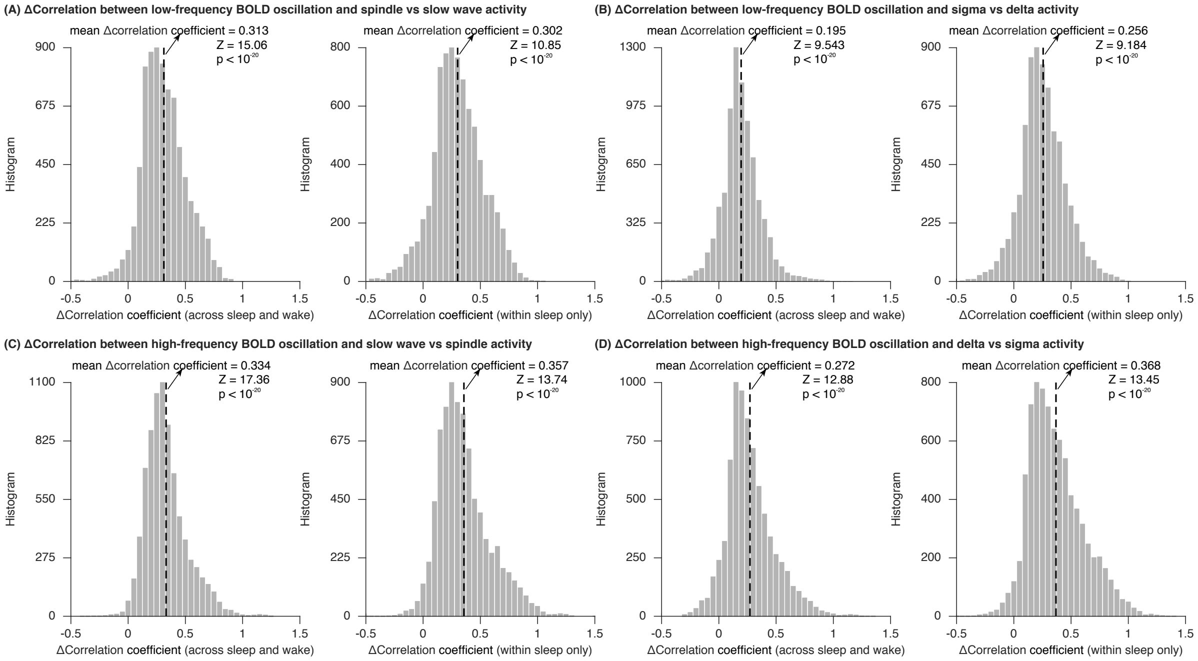


**Figure S2. Selective correlation between BOLD oscillations and sleep spindles versus slow waves.** The time course of low-frequency or high-frequency BOLD oscillation power was correlated against the time course of spindle or sigma activity versus the time course of slow wave or delta activity, and the difference in correlation coefficient was calculated, on a participant-by-participant, ROI-by-ROI basis, across sleep and wake (N = 1500 time points), or within sleep only (N = 1000 time points). The distributions of difference in correlation coefficient across all participants and ROIs were plotted, where each value reflected the result from a single participant and a single ROI. Low-frequency BOLD oscillation correlated significantly stronger with spindle or sigma activity than with slow wave or delta activity ($T > 9.183$, $p < 10^{-20}$). By contrast, high-frequency BOLD oscillation correlated significantly stronger with slow wave or delta activity than with spindle or sigma activity ($T > 12.878$, $p < 10^{-20}$). The figure shows the average results of two hemispheres across participants S1 to S36.

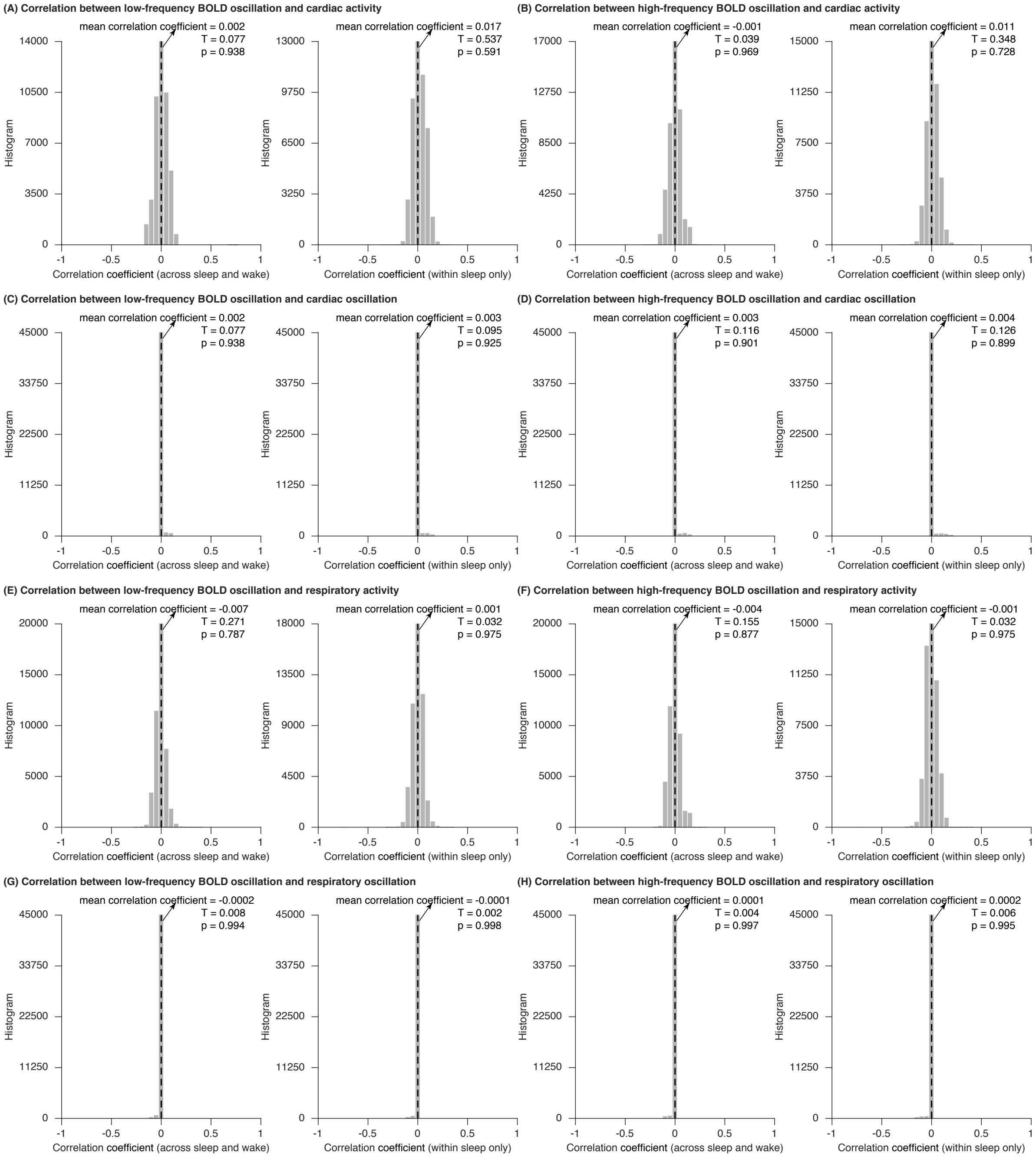


**Figure S3. Relation between BOLD oscillations and cardiac or respiratory activity.** The time course of low-frequency or high-frequency BOLD oscillation power was correlated against the time course of cardiac or respiratory activity (derived from respiration and pulse oximetry data) or the time course of cardiac or respiratory oscillation power (derived from FFT analysis to cardiac and respiratory time series), on a participant-by-participant, ROI-by-ROI basis, across sleep and wake (N = 1500 time points), or within sleep only (N = 1000 time points). The distributions of correlation coefficient across all participants and ROIs were plotted, where each value reflected the result from a single participant and a single ROI. At a data length of 1500 and 1000 time points, the threshold correlation coefficient for establishing statistical significance are 0.051 and 0.062 (before correction for multiple comparisons) or 0.136 and 0.166 (after Bonferroni correction for 374976 comparisons), respectively. The correlations between BOLD oscillation power and cardiac or respiratory regressors were much smaller than the threshold correlation coefficient, and corresponded to $T < 0.233$, $p > 0.815$. The figure shows the average results of two hemispheres across participants S1 to S36.

**Table 1. Sleep architecture of participants**

| Participants with Sustained Epochs of N2 or N3 Sleep | | | | | |
|---|---|---|---|---|---|
| **Participant** | **%N1** | **%N2** | **%N3** | **%N1/2/3** | **%N2/3** |
| **S1** | 19.47 | 18.20 | 48.53 | 86.20 | 66.73 |
| **S2** | 20.20 | 28.33 | 35.53 | 84.07 | 63.87 |
| **S3** | 35.60 | 21.13 | 0.00 | 56.73 | 21.13 |
| **S4** | 22.81 | 29.30 | 5.82 | 57.93 | 35.12 |
| **S5** | 6.73 | 26.00 | 0.00 | 32.73 | 26.00 |
| **S6** | 14.40 | 24.47 | 0.00 | 38.87 | 24.47 |
| **S7** | 27.67 | 39.20 | 0.00 | 66.87 | 39.20 |
| **S8** | 46.53 | 16.33 | 0.00 | 62.87 | 16.33 |
| **S9** | 8.73 | 28.80 | 0.00 | 37.53 | 28.80 |
| **S10** | 31.84 | 10.57 | 0.00 | 42.41 | 10.57 |
| **S11** | 13.93 | 44.13 | 26.00 | 84.07 | 70.13 |
| **S12** | 28.80 | 46.13 | 0.00 | 74.93 | 46.13 |
| **S13** | 27.89 | 23.21 | 0.00 | 51.10 | 23.21 |
| **S14** | 5.87 | 41.33 | 45.13 | 92.33 | 86.47 |
| **S15** | 20.20 | 39.33 | 0.00 | 59.53 | 39.33 |
| **S16** | 21.33 | 46.47 | 10.53 | 78.33 | 57.00 |
| **S17** | 18.27 | 23.20 | 42.60 | 84.07 | 65.80 |
| **S18** | 14.47 | 23.47 | 38.53 | 76.47 | 62.00 |
| **S19** | 13.47 | 32.67 | 14.87 | 61.00 | 47.53 |
| **S20** | 13.47 | 43.80 | 33.60 | 90.87 | 77.40 |
| **S21** | 21.13 | 35.00 | 27.93 | 84.07 | 62.93 |
| **S22** | 14.27 | 9.60 | 25.00 | 48.87 | 34.60 |
| **S23** | 22.33 | 20.00 | 0.00 | 42.33 | 20.00 |
| **S24** | 34.67 | 33.13 | 0.00 | 67.80 | 33.13 |
| **S25** | 18.13 | 26.87 | 24.07 | 69.07 | 50.93 |
| **S26** | 30.53 | 14.47 | 0.00 | 45.00 | 14.47 |
| **S27** | 36.40 | 55.87 | 0.00 | 92.27 | 55.87 |
| **S28** | 21.67 | 33.80 | 8.00 | 63.47 | 41.80 |
| **S29** | 6.73 | 19.27 | 66.73 | 92.73 | 86.00 |
| **S30** | 13.60 | 29.73 | 0.00 | 43.33 | 29.73 |
| **S31** | 23.14 | 13.51 | 39.53 | 76.19 | 53.04 |
| **S32** | 23.87 | 14.47 | 0.00 | 38.33 | 14.47 |
| **S33** | 17.67 | 19.27 | 28.87 | 65.80 | 48.13 |
| **S34** | 7.73 | 19.27 | 50.93 | 77.93 | 70.20 |
| **S35** | 31.73 | 55.27 | 0.00 | 87.00 | 55.27 |
| **S36** | 18.20 | 63.93 | 0.00 | 82.13 | 63.93 |
| **Participants without Sustained Epochs of N2 or N3 Sleep** | | | | | |
| **Participant** | **%N1** | **%N2** | **%N3** | **%N1/2/3** | **%N2/3** |
| **S37** | 1.93 | 0.00 | 0.00 | 1.93 | 0.00 |
| **S38** | 51.87 | 7.73 | 0.00 | 59.60 | 7.73 |
| **S39** | 90.87 | 0.00 | 0.00 | 90.87 | 0.00 |
| **S40** | 3.87 | 0.00 | 0.00 | 3.87 | 0.00 |
| **S41** | 14.40 | 0.00 | 0.00 | 14.40 | 0.00 |
| **S42** | 93.67 | 0.00 | 0.00 | 93.67 | 0.00 |
| **S43** | 6.27 | 0.00 | 0.00 | 6.27 | 0.00 |
| **S44** | 50.53 | 0.00 | 0.00 | 50.53 | 0.00 |
| **S45** | 14.87 | 0.00 | 0.00 | 14.87 | 0.00 |
| **S46** | 2.87 | 0.00 | 0.00 | 2.87 | 0.00 |
| **S47** | 22.34 | 0.00 | 0.00 | 22.34 | 0.00 |
| **S48** | 5.87 | 0.00 | 0.00 | 5.87 | 0.00 |
| **S49** | 5.67 | 0.00 | 0.00 | 5.67 | 0.00 |
| **S50** | 0.00 | 0.00 | 0.00 | 0.00 | 0.00 |
| **S51** | 43.07 | 4.80 | 0.00 | 47.87 | 4.80 |
| **S52** | 1.87 | 0.00 | 0.00 | 1.87 | 0.00 |
| **S53** | 25.00 | 0.00 | 0.00 | 25.00 | 0.00 |
| **S54** | 20.67 | 0.00 | 0.00 | 20.67 | 0.00 |
| **S55** | 68.20 | 4.80 | 0.00 | 73.00 | 4.80 |
| **S56** | 7.82 | 0.00 | 0.00 | 7.82 | 0.00 |
| **S57** | 20.60 | 3.87 | 0.00 | 24.47 | 3.87 |
| **S58** | 13.53 | 0.00 | 0.00 | 13.53 | 0.00 |

**Table S1. Sleep architecture of participants.** Simultaneous fMRI and polysomnographic EEG recordings were acquired from 58 non-sleep-deprived participants. Recordings started at around 20:00 and lasted for 50 minutes. 36 out of 58 participants displaying sustained epochs of N2 and N3 sleep were included in subsequent analyses. Listed in the table are the % time spent in N1, N2, N3 sleep or their combinations.

**Table S2. Regional distribution of low-frequency BOLD oscillation**

| Fine atlas | | | |
|---|---|---|---|
| Cortex | Power | | Frequency |
| | % Increase | T Statistics | Hz |
| **25** | 405.39 | 9.983 | 0.0570 |
| **V6** | 348.36 | 14.679 | 0.0522 |
| **V1** | 303.87 | 13.854 | 0.0523 |
| **52** | 299.65 | 12.808 | 0.0555 |
| **DVT** | 294.37 | 13.405 | 0.0530 |
| **MBelt** | 292.60 | 12.560 | 0.0549 |
| **LBelt** | 281.82 | 13.261 | 0.0533 |
| **V6A** | 272.95 | 13.298 | 0.0533 |
| **PBelt** | 269.18 | 13.105 | 0.0530 |
| **V2** | 265.80 | 13.274 | 0.0531 |
| **POS2** | 260.26 | 13.327 | 0.0527 |
| **A1** | 259.49 | 13.425 | 0.0533 |
| **ProS** | 256.86 | 13.503 | 0.0542 |
| **POS1** | 251.73 | 12.997 | 0.0539 |
| **7m** | 250.16 | 12.944 | 0.0532 |
| **MT** | 248.19 | 12.838 | 0.0540 |
| **31pd** | 247.57 | 12.231 | 0.0539 |
| **V3A** | 243.99 | 12.764 | 0.0536 |
| **VMV1** | 240.37 | 13.048 | 0.0543 |
| **a24** | 240.16 | 9.844 | 0.0560 |
| **IPS1** | 237.73 | 13.365 | 0.0534 |
| **V7** | 237.33 | 12.376 | 0.0536 |
| **IP0** | 234.35 | 12.340 | 0.0540 |
| **TPOJ1** | 233.18 | 12.563 | 0.0546 |
| **7PL** | 232.79 | 12.670 | 0.0524 |
| **MST** | 231.27 | 12.601 | 0.0540 |
| **V3** | 230.39 | 12.552 | 0.0538 |
| **RI** | 230.15 | 12.706 | 0.0548 |
| **A4** | 229.80 | 12.397 | 0.0528 |
| **7Pm** | 228.02 | 13.058 | 0.0529 |
| **V8** | 224.71 | 12.679 | 0.0544 |
| **TPOJ2** | 222.22 | 11.976 | 0.0543 |
| **VMV3** | 220.83 | 12.600 | 0.0544 |
| **VIP** | 220.59 | 12.155 | 0.0546 |
| **LO3** | 220.01 | 12.430 | 0.0542 |
| **v23ab** | 219.60 | 13.082 | 0.0532 |
| **7Am** | 217.92 | 11.925 | 0.0538 |
| **OP1** | 217.45 | 11.124 | 0.0561 |
| **VVC** | 216.56 | 11.429 | 0.0557 |
| **V3B** | 215.59 | 12.806 | 0.0539 |
| **MIP** | 211.34 | 12.160 | 0.0538 |
| **RSC** | 209.76 | 11.662 | 0.0551 |
| **PIT** | 209.68 | 11.886 | 0.0542 |
| **TPOJ3** | 209.37 | 12.649 | 0.0544 |
| **LO1** | 209.04 | 12.193 | 0.0545 |
| **3b** | 208.65 | 11.552 | 0.0542 |
| **OP2-3** | 208.21 | 10.686 | 0.0574 |
| **PCV** | 207.92 | 11.756 | 0.0542 |
| **STSvp** | 206.98 | 11.434 | 0.0542 |
| **3a** | 206.80 | 11.729 | 0.0549 |
| **23c** | 206.52 | 12.044 | 0.0543 |
| **V4** | 206.10 | 11.857 | 0.0545 |
| **V3CD** | 206.07 | 11.832 | 0.0546 |

| | | | |
|---|---|---|---|
| **p24pr** | 206.00 | 12.015 | 0.0549 |
| **Ig** | 205.92 | 11.886 | 0.0561 |
| **10r** | 205.36 | 9.246 | 0.0553 |
| **STV** | 204.67 | 12.094 | 0.0548 |
| **PI** | 204.33 | 11.099 | 0.0567 |
| **VMV2** | 204.15 | 12.496 | 0.0551 |
| **PSL** | 203.79 | 11.659 | 0.0544 |
| **TA2** | 203.65 | 11.833 | 0.0543 |
| **31a** | 202.98 | 12.150 | 0.0538 |
| **PreS** | 202.94 | 11.928 | 0.0556 |
| **6d** | 201.81 | 11.241 | 0.0556 |
| **IFJa** | 201.60 | 10.210 | 0.0553 |
| **31pv** | 201.53 | 11.220 | 0.0540 |
| **FFC** | 201.32 | 11.854 | 0.0551 |
| **47m** | 198.54 | 10.175 | 0.0563 |
| **24dd** | 198.31 | 11.206 | 0.0561 |
| **MI** | 198.12 | 11.248 | 0.0559 |
| **5m** | 196.69 | 11.790 | 0.0563 |
| **PoI1** | 196.65 | 11.007 | 0.0568 |
| **a24pr** | 193.33 | 11.143 | 0.0557 |
| **s32** | 191.87 | 9.917 | 0.0561 |
| **24dv** | 191.25 | 11.019 | 0.0560 |
| **PGp** | 190.45 | 11.290 | 0.0549 |
| **PFt** | 189.10 | 11.561 | 0.0545 |
| **IP1** | 189.06 | 11.315 | 0.0541 |
| **d23ab** | 189.00 | 11.695 | 0.0539 |
| **PHA1** | 188.49 | 10.964 | 0.0566 |
| **FST** | 188.11 | 11.447 | 0.0548 |
| **LIPd** | 188.07 | 11.481 | 0.0544 |
| **IP2** | 188.02 | 10.754 | 0.0548 |
| **STGa** | 187.04 | 10.971 | 0.0556 |
| **AAIC** | 187.00 | 10.387 | 0.0567 |
| **5L** | 186.89 | 10.881 | 0.0560 |
| **7AL** | 186.65 | 11.579 | 0.0552 |
| **6r** | 185.63 | 10.188 | 0.0553 |
| **A5** | 185.04 | 11.568 | 0.0556 |
| **10pp** | 185.03 | 8.321 | 0.0560 |
| **PH** | 184.97 | 11.060 | 0.0554 |
| **PFcm** | 184.93 | 10.868 | 0.0561 |
| **PHT** | 184.93 | 10.888 | 0.0551 |
| **LIPv** | 184.62 | 11.868 | 0.0545 |
| **EC** | 184.01 | 9.676 | 0.0579 |
| **1** | 183.51 | 10.859 | 0.0555 |
| **STSdp** | 182.55 | 10.774 | 0.0567 |
| **FOP2** | 181.86 | 10.694 | 0.0577 |
| **STSva** | 181.03 | 10.832 | 0.0558 |
| **V4t** | 180.84 | 12.109 | 0.0548 |
| **Pir** | 179.56 | 10.190 | 0.0573 |
| **7PC** | 178.47 | 11.492 | 0.0545 |
| **PEF** | 178.45 | 11.377 | 0.0555 |
| **6v** | 178.12 | 10.805 | 0.0556 |
| **47s** | 178.10 | 10.351 | 0.0563 |
| **LO2** | 176.86 | 11.422 | 0.0547 |
| **AIP** | 176.73 | 11.203 | 0.0548 |
| **PGi** | 176.30 | 10.843 | 0.0550 |
| **H** | 175.78 | 10.588 | 0.0565 |
| **23d** | 175.75 | 11.150 | 0.0548 |
| **IFJp** | 175.67 | 11.205 | 0.0550 |

| | | | |
|---|---|---|---|
| **PHA2** | 175.13 | 11.015 | 0.0573 |
| **47l** | 174.16 | 10.077 | 0.0554 |
| **5mv** | 173.33 | 11.255 | 0.0555 |
| **STSda** | 172.83 | 10.336 | 0.0570 |
| **FOP3** | 172.55 | 10.581 | 0.0575 |
| **33pr** | 172.31 | 11.371 | 0.0565 |
| **PoI2** | 171.75 | 11.168 | 0.0562 |
| **43** | 170.93 | 10.634 | 0.0566 |
| **IFSp** | 170.69 | 10.360 | 0.0549 |
| **TGd** | 170.57 | 9.423 | 0.0564 |
| **OP4** | 170.53 | 10.838 | 0.0563 |
| **4** | 170.52 | 10.602 | 0.0565 |
| **FEF** | 170.20 | 10.296 | 0.0560 |
| **TE1p** | 169.66 | 10.194 | 0.0560 |
| **10v** | 169.47 | 8.941 | 0.0551 |
| **6mp** | 169.14 | 10.477 | 0.0563 |
| **p24** | 168.63 | 10.177 | 0.0560 |
| **TE1m** | 168.14 | 10.046 | 0.0572 |
| **p32pr** | 167.87 | 10.145 | 0.0570 |
| **AVI** | 167.19 | 10.231 | 0.0566 |
| **PF** | 166.92 | 10.365 | 0.0556 |
| **PHA3** | 166.62 | 11.068 | 0.0562 |
| **2** | 165.14 | 10.730 | 0.0558 |
| **PFop** | 162.75 | 11.183 | 0.0557 |
| **45** | 162.54 | 10.034 | 0.0561 |
| **55b** | 161.30 | 10.566 | 0.0558 |
| **SCEF** | 159.78 | 9.836 | 0.0567 |
| **p47r** | 159.69 | 9.384 | 0.0556 |
| **PFm** | 159.46 | 10.360 | 0.0550 |
| **PGs** | 156.93 | 10.111 | 0.0549 |
| **44** | 156.61 | 10.195 | 0.0559 |
| **6a** | 155.24 | 9.768 | 0.0564 |
| **10d** | 154.62 | 8.816 | 0.0543 |
| **SFL** | 154.55 | 8.614 | 0.0556 |
| **TE2p** | 154.23 | 9.777 | 0.0574 |
| **a10p** | 153.41 | 9.288 | 0.0550 |
| **a9-46v** | 153.10 | 9.030 | 0.0552 |
| **FOP1** | 152.71 | 10.086 | 0.0577 |
| **pOFC** | 152.17 | 9.433 | 0.0577 |
| **13l** | 150.69 | 9.471 | 0.0569 |
| **FOP5** | 150.68 | 9.791 | 0.0581 |
| **6ma** | 150.50 | 8.875 | 0.0565 |
| **a47r** | 150.46 | 8.945 | 0.0558 |
| **9m** | 150.27 | 8.737 | 0.0558 |
| **a32pr** | 150.15 | 9.685 | 0.0573 |
| **OFC** | 150.12 | 9.093 | 0.0572 |
| **PeEc** | 149.24 | 9.134 | 0.0582 |
| **TE1a** | 148.68 | 9.585 | 0.0565 |
| **8Ad** | 147.48 | 9.097 | 0.0564 |
| **FOP4** | 146.79 | 9.937 | 0.0575 |
| **11l** | 146.54 | 8.893 | 0.0568 |
| **p9-46v** | 146.14 | 9.709 | 0.0559 |
| **TGv** | 144.63 | 9.281 | 0.0580 |
| **s6-8** | 143.70 | 9.514 | 0.0565 |
| **8BL** | 143.65 | 7.804 | 0.0558 |
| **46** | 143.25 | 9.154 | 0.0555 |
| **p32** | 142.76 | 9.518 | 0.0562 |
| **IFSa** | 142.57 | 9.595 | 0.0551 |

| | | | |
|---|---|---|---|
| **9p** | 142.53 | 8.629 | 0.0557 |
| **9-46d** | 141.47 | 9.040 | 0.0557 |
| **8Av** | 141.16 | 8.626 | 0.0564 |
| **8C** | 141.04 | 9.495 | 0.0567 |
| **p10p** | 140.07 | 8.701 | 0.0550 |
| **TF** | 140.04 | 9.390 | 0.0581 |
| **9a** | 139.42 | 8.609 | 0.0552 |
| **TE2a** | 138.42 | 9.062 | 0.0582 |
| **i6-8** | 136.11 | 8.699 | 0.0562 |
| **8BM** | 133.16 | 8.882 | 0.0569 |
| **d32** | 132.63 | 9.412 | 0.0569 |
| **Subcortex** | **Power** | | **Frequency** |
| | **% Increase** | **T Statistics** | **Hz** |
| **Hypothalamus** | 217.46 | 9.774 | 0.0572 |
| **Thalamus (Anterior)** | 215.18 | 10.412 | 0.0568 |
| **Thalamus (Parafascicular)** | 211.07 | 10.925 | 0.0578 |
| **Thalamus (Medial Dorsal)** | 203.08 | 10.447 | 0.0579 |
| **Thalamus (Lateral Dorsal)** | 198.81 | 10.756 | 0.0579 |
| **Basal Forebrain** | 191.58 | 9.572 | 0.0573 |
| **Thalamus (Pulvinar)** | 190.12 | 10.530 | 0.0578 |
| **Cerebellum (HX)** | 187.77 | 10.334 | 0.0577 |
| **Cerebellum (HIIV)** | 184.38 | 9.953 | 0.0585 |
| **Cerebellum (HV)** | 181.73 | 9.366 | 0.0582 |
| **Midbrain** | 179.24 | 10.313 | 0.0577 |
| **Cerebellum (HVIIIb)** | 177.49 | 9.397 | 0.0573 |
| **Thalamus (Ventral Posterior)** | 175.30 | 10.237 | 0.0581 |
| **Thalamus (Ventral Lateral)** | 174.20 | 9.879 | 0.0581 |
| **Striatum (Parietal Network)** | 173.78 | 10.068 | 0.0581 |
| **Cerebellum (HIX)** | 167.01 | 9.340 | 0.0581 |
| **Medulla** | 165.14 | 9.255 | 0.0587 |
| **Amygdala** | 164.15 | 9.658 | 0.0572 |
| **Cerebellum (HVIIIa)** | 163.70 | 9.096 | 0.0573 |
| **Pons** | 163.19 | 9.043 | 0.0585 |
| **Thalamus (Medial Geniculate)** | 161.50 | 9.926 | 0.0566 |
| **Striatum (Central Network)** | 161.48 | 9.837 | 0.0585 |
| **Thalamus (Ventral Anterior)** | 160.88 | 10.098 | 0.0576 |
| **Hippocampus** | 160.11 | 9.735 | 0.0576 |
| **Thalamus (Lateral Posterior)** | 151.83 | 9.988 | 0.0586 |
| **Cerebellum (HVIIb)** | 150.45 | 8.738 | 0.0581 |
| **Cerebellum (Dentate)** | 147.12 | 9.308 | 0.0579 |
| **Thalamus (Lateral Geniculate)** | 146.85 | 8.902 | 0.0575 |
| **Cerebellum (HVI)** | 146.11 | 8.981 | 0.0580 |
| **Striatum (Frontal Network)** | 145.93 | 9.278 | 0.0581 |
| **Striatum (Limbic Network)** | 144.16 | 9.439 | 0.0575 |
| **Cerebellum (CrusI)** | 141.40 | 8.521 | 0.0580 |
| **Cerebellum (CrusII)** | 140.45 | 8.117 | 0.0584 |
| **Thalamus (Centromedian)** | 137.37 | 9.587 | 0.0574 |
| **Striatum (Temporal Network)** | 125.65 | 8.778 | 0.0579 |
| **Striatum (Occipital Network)** | 123.60 | 8.851 | 0.0586 |
| **Coarse atlas** | | | |
| **Cortex** | **Power** | | **Frequency** |
| | **% Increase** | **T Statistics** | **Hz** |
| **Pericalcarine (V1)** | 338.03 | 14.460 | 0.0526 |

| | | | |
|---|---|---|---|
| **Cuneus (V2)** | 312.90 | 14.392 | 0.0522 |
| **Transverse Temporal (A1)** | 278.44 | 12.814 | 0.0538 |
| **Lingual (V2)** | 259.42 | 13.497 | 0.0531 |
| **Precuneus** | 220.46 | 12.227 | 0.0534 |
| **Isthmus Cingulate** | 208.30 | 11.699 | 0.0540 |
| **Rostral Anterior Cingulate** | 205.85 | 9.543 | 0.0564 |
| **Superior Parietal** | 199.61 | 11.657 | 0.0538 |
| **Superior Temporal Sulcus** | 199.15 | 11.192 | 0.0547 |
| **Medial Orbitofrontal** | 197.48 | 8.971 | 0.0562 |
| **Superior Temporal** | 192.18 | 11.104 | 0.0550 |
| **Lateral Occipital** | 190.52 | 11.357 | 0.0544 |
| **Paracentral** | 180.25 | 11.018 | 0.0561 |
| **Parahippocampal** | 180.21 | 10.389 | 0.0572 |
| **Posterior Cingulate** | 177.62 | 11.196 | 0.0548 |
| **Postcentral (S1)** | 177.13 | 10.782 | 0.0554 |
| **Caudal Anterior Cingulate** | 177.04 | 10.610 | 0.0561 |
| **Fusiform** | 174.27 | 10.655 | 0.0561 |
| **Inferior Parietal** | 172.51 | 10.679 | 0.0547 |
| **Insula** | 171.75 | 10.550 | 0.0565 |
| **Supra Marginal** | 170.69 | 10.652 | 0.0554 |
| **Middle Temporal** | 167.72 | 10.034 | 0.0558 |
| **Precentral** | 163.87 | 10.323 | 0.0560 |
| **Pars Opercularis** | 154.00 | 9.683 | 0.0559 |
| **Entorhinal** | 153.56 | 9.028 | 0.0581 |
| **Lateral Orbitofrontal** | 148.36 | 8.987 | 0.0569 |
| **Pars Triangularis** | 146.17 | 9.546 | 0.0559 |
| **Inferior Temporal** | 144.65 | 9.482 | 0.0572 |
| **Pars Orbitalis** | 143.90 | 8.995 | 0.0560 |
| **Rostral Middle Frontal** | 141.55 | 8.720 | 0.0555 |
| **Superior Frontal** | 136.01 | 8.496 | 0.0562 |
| **Caudal Middle Frontal** | 133.74 | 8.697 | 0.0566 |
| **Subcortex** | **Power** | | **Frequency** |
| | **% Increase** | **T Statistics** | **Hz** |
| **Hypothalamus** | 217.46 | 9.774 | 0.0572 |
| **Basal Forebrain** | 191.58 | 9.572 | 0.0573 |
| **Striatum** | 186.52 | 10.716 | 0.0576 |
| **Thalamus** | 183.27 | 10.874 | 0.0575 |
| **Midbrain** | 179.24 | 10.313 | 0.0577 |
| **Medulla** | 165.14 | 9.255 | 0.0587 |
| **Amygdala** | 164.15 | 9.658 | 0.0572 |
| **Pons** | 163.19 | 9.043 | 0.0585 |
| **Hippocampus** | 160.11 | 9.735 | 0.0576 |
| **Cerebellum** | 149.40 | 8.715 | 0.0580 |

**Table S2. Regional distributions of low-frequency BOLD oscillation.** BOLD power spectrogram was acquired on a voxel-level and the power spectrogram of each ROI was calculated as the normalized average across all voxels within this ROI. Based on the power spectrogram, we identified the spectral peaks in lower-frequency and higher-frequency ranges, traced the time courses of oscillation power, calculated the increase in oscillation power from wake to sleep, and the statistical difference (T value) in oscillation power between wake and sleep. Listed in the table are the spectral peak (Hz) and oscillation power (% increase from wake to sleep, statistical difference between wake and sleep) of low-frequency BOLD oscillation, averaged across two hemispheres and participants S1 to S36.

**Table S3. Regional distribution of high-frequency BOLD oscillation**

| Fine atlas | | | |
|---|---|---|---|
| **Cortex** | **Power** | | **Frequency** |
| | **% Increase** | **T Statistics** | **Hz** |
| **25** | 344.03 | 11.8912 | 0.1730 |
| **PI** | 321.37 | 13.1483 | 0.1683 |
| **EC** | 303.17 | 12.4000 | 0.1738 |
| **Pir** | 297.46 | 11.8269 | 0.1704 |
| **PreS** | 297.38 | 12.9346 | 0.1690 |
| **10r** | 285.20 | 11.1225 | 0.1729 |
| **V6A** | 283.65 | 12.2329 | 0.1689 |
| **TA2** | 274.59 | 12.4598 | 0.1698 |
| **VMV2** | 274.08 | 12.2558 | 0.1723 |
| **47s** | 265.97 | 12.0338 | 0.1728 |
| **10v** | 263.88 | 11.2910 | 0.1745 |
| **TGd** | 261.67 | 11.5398 | 0.1745 |
| **PBelt** | 261.56 | 12.6764 | 0.1677 |
| **STGa** | 257.18 | 12.1507 | 0.1723 |
| **FOP1** | 254.38 | 11.8067 | 0.1705 |
| **PHA1** | 249.42 | 12.0579 | 0.1709 |
| **10pp** | 248.78 | 11.4472 | 0.1760 |
| **PHA2** | 246.60 | 11.9925 | 0.1713 |
| **Ig** | 246.05 | 11.4416 | 0.1694 |
| **VMV1** | 245.58 | 12.0403 | 0.1692 |
| **A4** | 245.29 | 12.7759 | 0.1687 |
| **6d** | 243.66 | 12.0341 | 0.1717 |
| **V8** | 243.60 | 11.9106 | 0.1692 |
| **a10p** | 242.97 | 11.3384 | 0.1747 |
| **10d** | 242.39 | 11.3141 | 0.1751 |
| **SFL** | 241.84 | 11.3300 | 0.1736 |
| **STSva** | 240.82 | 11.8870 | 0.1710 |
| **VVC** | 239.93 | 11.7786 | 0.1709 |
| **PHA3** | 239.61 | 11.4799 | 0.1716 |
| **7PL** | 239.38 | 11.7475 | 0.1693 |
| **VIP** | 237.49 | 11.5620 | 0.1693 |
| **A1** | 236.67 | 12.5086 | 0.1676 |
| **6ma** | 234.82 | 11.2418 | 0.1732 |
| **d32** | 234.52 | 11.1155 | 0.1713 |
| **p10p** | 234.23 | 11.4408 | 0.1744 |
| **PIT** | 233.66 | 11.7412 | 0.1698 |
| **V7** | 233.57 | 11.6141 | 0.1689 |
| **pOFC** | 233.53 | 11.1618 | 0.1743 |
| **PoI2** | 233.39 | 11.9871 | 0.1698 |
| **FFC** | 233.01 | 11.7779 | 0.1716 |
| **AAIC** | 231.84 | 11.5292 | 0.1704 |
| **p24** | 231.21 | 11.8338 | 0.1706 |
| **52** | 230.06 | 12.4282 | 0.1680 |
| **a24** | 230.01 | 11.2710 | 0.1710 |
| **PGp** | 225.92 | 11.2826 | 0.1704 |
| **V1** | 225.19 | 12.0635 | 0.1651 |
| **TE2p** | 225.06 | 12.7326 | 0.1728 |
| **LBelt** | 225.06 | 11.4890 | 0.1669 |
| **PoI1** | 224.99 | 11.8680 | 0.1690 |
| **7Pm** | 224.64 | 11.5834 | 0.1682 |
| **PGs** | 224.55 | 11.2974 | 0.1700 |
| **MBelt** | 222.81 | 12.5194 | 0.1680 |
| **8BL** | 222.42 | 11.1726 | 0.1761 |

| | | | |
|---|---|---|---|
| **STSda** | 222.35 | 11.7037 | 0.1710 |
| **TPOJ3** | 221.51 | 11.3535 | 0.1692 |
| **13l** | 220.92 | 10.9157 | 0.1729 |
| **a9-46v** | 220.67 | 11.3963 | 0.1739 |
| **1** | 220.50 | 11.5252 | 0.1716 |
| **PeEc** | 220.20 | 11.3965 | 0.1738 |
| **7Am** | 219.87 | 11.2666 | 0.1689 |
| **ProS** | 219.46 | 12.2738 | 0.1687 |
| **RSC** | 219.42 | 11.4177 | 0.1684 |
| **DVT** | 219.26 | 11.8262 | 0.1676 |
| **PSL** | 218.95 | 11.9705 | 0.1689 |
| **IP1** | 218.89 | 11.3250 | 0.1694 |
| **STV** | 218.65 | 11.5968 | 0.1701 |
| **p32** | 218.36 | 11.2104 | 0.1722 |
| **9-46d** | 218.31 | 11.2019 | 0.1731 |
| **23d** | 217.58 | 11.2992 | 0.1688 |
| **p47r** | 216.87 | 11.5804 | 0.1732 |
| **6mp** | 216.44 | 11.1454 | 0.1710 |
| **p24pr** | 216.25 | 11.7479 | 0.1690 |
| **IP0** | 214.58 | 11.3411 | 0.1685 |
| **FOP5** | 214.54 | 11.3581 | 0.1710 |
| **AVI** | 214.16 | 11.3859 | 0.1709 |
| **s32** | 214.03 | 11.1490 | 0.1730 |
| **TF** | 213.86 | 11.1954 | 0.1727 |
| **POS2** | 213.60 | 11.7008 | 0.1663 |
| **V3B** | 213.36 | 12.1153 | 0.1676 |
| **A5** | 213.04 | 11.8113 | 0.1707 |
| **8Av** | 212.71 | 11.2145 | 0.1740 |
| **TPOJ1** | 212.62 | 12.0461 | 0.1697 |
| **MI** | 212.32 | 11.7167 | 0.1700 |
| **POS1** | 211.74 | 12.0110 | 0.1676 |
| **IPS1** | 211.73 | 11.4875 | 0.1681 |
| **24dv** | 211.68 | 11.3154 | 0.1694 |
| **a47r** | 211.66 | 11.3538 | 0.1738 |
| **33pr** | 211.59 | 11.8635 | 0.1703 |
| **OP2-3** | 211.32 | 11.5286 | 0.1699 |
| **VMV3** | 211.16 | 12.3646 | 0.1688 |
| **5mv** | 211.05 | 11.2909 | 0.1689 |
| **i6-8** | 210.91 | 11.5346 | 0.1735 |
| **V3A** | 210.84 | 11.3521 | 0.1685 |
| **TE1p** | 210.65 | 11.6089 | 0.1714 |
| **V2** | 210.57 | 11.6881 | 0.1668 |
| **V6** | 209.19 | 11.8470 | 0.1655 |
| **TE1a** | 208.54 | 11.6521 | 0.1726 |
| **3b** | 208.32 | 11.3749 | 0.1692 |
| **TPOJ2** | 208.28 | 11.6333 | 0.1692 |
| **MST** | 208.03 | 11.8935 | 0.1682 |
| **47l** | 207.35 | 11.5884 | 0.1723 |
| **LO1** | 206.88 | 11.7990 | 0.1693 |
| **LO3** | 206.51 | 11.9749 | 0.1695 |
| **s6-8** | 206.33 | 11.8442 | 0.1738 |
| **STSdp** | 206.21 | 11.4933 | 0.1704 |
| **TGv** | 206.17 | 11.4353 | 0.1739 |
| **FOP2** | 206.02 | 11.8773 | 0.1710 |
| **31a** | 205.82 | 11.5237 | 0.1678 |
| **H** | 205.49 | 12.3485 | 0.1704 |
| **45** | 205.22 | 11.3101 | 0.1717 |
| **V3** | 205.14 | 11.3914 | 0.1680 |

| | | | |
|---|---|---|---|
| **LO2** | 205.14 | 11.5277 | 0.1698 |
| **9a** | 204.88 | 11.1462 | 0.1749 |
| **PHT** | 204.88 | 11.2728 | 0.1707 |
| **p32pr** | 204.22 | 11.2789 | 0.1704 |
| **V4** | 204.09 | 11.4711 | 0.1692 |
| **31pd** | 204.06 | 11.6400 | 0.1676 |
| **47m** | 204.01 | 11.6015 | 0.1718 |
| **OFC** | 203.97 | 11.4106 | 0.1735 |
| **7m** | 203.92 | 11.7779 | 0.1674 |
| **7AL** | 203.01 | 11.2544 | 0.1699 |
| **STSvp** | 202.66 | 11.5435 | 0.1700 |
| **AIP** | 201.52 | 11.4222 | 0.1692 |
| **31pv** | 201.50 | 11.5633 | 0.1681 |
| **RI** | 200.45 | 11.8961 | 0.1680 |
| **PFcm** | 200.38 | 11.5955 | 0.1699 |
| **PH** | 200.12 | 11.5250 | 0.1709 |
| **V3CD** | 199.02 | 11.3479 | 0.1692 |
| **p9-46v** | 198.36 | 11.0261 | 0.1728 |
| **PGi** | 198.33 | 11.1157 | 0.1694 |
| **24dd** | 198.11 | 11.1354 | 0.1694 |
| **TE1m** | 198.08 | 11.3872 | 0.1717 |
| **TE2a** | 197.86 | 11.1450 | 0.1724 |
| **MT** | 197.09 | 11.5017 | 0.1689 |
| **44** | 196.85 | 11.3204 | 0.1711 |
| **a24pr** | 196.77 | 11.5053 | 0.1694 |
| **v23ab** | 196.48 | 11.9916 | 0.1674 |
| **55b** | 196.32 | 11.5999 | 0.1712 |
| **7PC** | 196.29 | 11.2828 | 0.1705 |
| **6v** | 196.03 | 11.0759 | 0.1706 |
| **PEF** | 195.90 | 11.1990 | 0.1702 |
| **4** | 195.14 | 11.0332 | 0.1702 |
| **PFt** | 195.11 | 11.8055 | 0.1701 |
| **11l** | 194.69 | 11.1421 | 0.1734 |
| **8Ad** | 194.47 | 11.0944 | 0.1731 |
| **MIP** | 194.35 | 11.2122 | 0.1682 |
| **9p** | 194.21 | 11.0981 | 0.1744 |
| **9m** | 194.08 | 10.8206 | 0.1733 |
| **46** | 193.29 | 10.8964 | 0.1732 |
| **IFJp** | 192.87 | 11.8451 | 0.1702 |
| **2** | 192.72 | 11.3017 | 0.1701 |
| **6r** | 192.66 | 11.2014 | 0.1702 |
| **LIPv** | 192.62 | 11.4557 | 0.1695 |
| **PCV** | 191.02 | 11.4175 | 0.1678 |
| **d23ab** | 190.52 | 11.6778 | 0.1680 |
| **OP4** | 190.29 | 11.3430 | 0.1701 |
| **FST** | 189.52 | 11.5585 | 0.1693 |
| **PFm** | 189.49 | 11.1967 | 0.1700 |
| **43** | 188.11 | 11.4582 | 0.1702 |
| **FOP3** | 188.07 | 11.7135 | 0.1715 |
| **23c** | 188.01 | 11.1665 | 0.1682 |
| **IFSa** | 187.27 | 11.1631 | 0.1720 |
| **IFJa** | 186.54 | 11.3607 | 0.1707 |
| **IFSp** | 186.50 | 11.2415 | 0.1711 |
| **FOP4** | 186.46 | 11.5456 | 0.1711 |
| **PFop** | 186.26 | 11.6663 | 0.1705 |
| **3a** | 184.70 | 11.6747 | 0.1690 |
| **5m** | 184.45 | 11.0378 | 0.1696 |
| **PF** | 181.65 | 11.4194 | 0.1703 |

| | | | |
|---|---|---|---|
| **8C** | 181.29 | 10.7894 | 0.1718 |
| **OP1** | 180.58 | 11.4355 | 0.1700 |
| **LIPd** | 180.27 | 11.0688 | 0.1683 |
| **V4t** | 180.15 | 11.7202 | 0.1695 |
| **5L** | 178.26 | 11.2967 | 0.1702 |
| **FEF** | 177.72 | 11.2246 | 0.1711 |
| **IP2** | 177.71 | 11.0644 | 0.1694 |
| **8BM** | 175.74 | 11.0164 | 0.1711 |
| **a32pr** | 173.91 | 11.2006 | 0.1709 |
| **6a** | 170.90 | 10.8008 | 0.1705 |
| **SCEF** | 168.47 | 10.8592 | 0.1703 |
| **Subcortex** | **Power** | | **Frequency** |
| | **% Increase** | **T Statistics** | **Hz** |
| **Thalamus (Parafascicular)** | 428.23 | 12.3498 | 0.1728 |
| **Thalamus (Lateral Dorsal)** | 365.21 | 11.4822 | 0.1706 |
| **Thalamus (Anterior)** | 341.62 | 12.0021 | 0.1693 |
| **Hypothalamus** | 340.18 | 11.7229 | 0.1733 |
| **Cerebellum (HX)** | 328.31 | 12.2171 | 0.1729 |
| **Thalamus (Medial Dorsal)** | 306.96 | 11.8852 | 0.1712 |
| **Thalamus (Medial Geniculate)** | 293.95 | 12.1211 | 0.1696 |
| **Cerebellum (HIX)** | 284.69 | 12.0195 | 0.1752 |
| **Basal Forebrain** | 268.41 | 12.0110 | 0.1732 |
| **Cerebellum (HVIIIa)** | 266.21 | 11.5614 | 0.1762 |
| **Amygdala** | 265.19 | 11.5314 | 0.1728 |
| **Cerebellum (HVIIIb)** | 258.40 | 11.9961 | 0.1760 |
| **Thalamus (Ventral Posterior)** | 251.03 | 11.5005 | 0.1711 |
| **Medulla** | 239.55 | 11.0568 | 0.1715 |
| **Midbrain** | 239.41 | 11.8140 | 0.1717 |
| **Thalamus (Pulvinar)** | 236.16 | 11.5566 | 0.1709 |
| **Hippocampus** | 236.10 | 11.3878 | 0.1722 |
| **Cerebellum (HIIV)** | 236.06 | 11.3452 | 0.1730 |
| **Cerebellum (HVIIb)** | 233.26 | 11.1379 | 0.1752 |
| **Thalamus (Ventral Lateral)** | 231.36 | 11.3785 | 0.1712 |
| **Thalamus (Centromedian)** | 227.77 | 11.4987 | 0.1717 |
| **Thalamus (Lateral Posterior)** | 227.66 | 10.9321 | 0.1704 |
| **Striatum (Limbic Network)** | 226.00 | 11.2582 | 0.1736 |
| **Cerebellum (HVI)** | 224.30 | 10.9621 | 0.1725 |
| **Pons** | 222.48 | 10.8429 | 0.1723 |
| **Thalamus (Ventral Anterior)** | 221.00 | 11.6988 | 0.1718 |
| **Cerebellum (HV)** | 215.43 | 11.0370 | 0.1726 |
| **Cerebellum (CrusII)** | 214.12 | 10.6736 | 0.1735 |
| **Cerebellum (CrusI)** | 213.92 | 10.6474 | 0.1731 |
| **Cerebellum (Dentate)** | 213.54 | 11.1972 | 0.1751 |
| **Striatum (Frontal Network)** | 208.56 | 11.1691 | 0.1719 |
| **Striatum (Parietal Network)** | 207.04 | 11.1188 | 0.1702 |
| **Thalamus (Lateral Geniculate)** | 202.26 | 11.3764 | 0.1723 |
| **Striatum (Occipital Network)** | 202.16 | 9.8077 | 0.1717 |
| **Striatum (Central Network)** | 192.40 | 11.0254 | 0.1695 |
| **Striatum (Temporal Network)** | 186.65 | 10.8316 | 0.1733 |
| **Coarse atlas** | | | |
| **Cortex** | **Power** | | **Frequency** |

| | % Increase | T Statistics | Hz |
|---|---|---|---|
| **Parahippocampal** | 247.85 | 11.8621 | 0.1715 |
| **Entorhinal** | 245.86 | 11.4860 | 0.1745 |
| **Medial Orbitofrontal** | 244.46 | 10.9848 | 0.1734 |
| **Superior Temporal** | 237.45 | 11.7668 | 0.1699 |
| **Rostral Anterior Cingulate** | 231.50 | 11.3533 | 0.1713 |
| **Pericalcarine (V1)** | 228.08 | 12.4096 | 0.1640 |
| **Cuneus (V2)** | 226.34 | 12.0146 | 0.1652 |
| **Transverse Temporal (A1)** | 224.88 | 12.5212 | 0.1679 |
| **Insula** | 217.90 | 11.4613 | 0.1699 |
| **Lingual (V2)** | 217.30 | 11.8974 | 0.1670 |
| **Pars Orbitalis** | 211.82 | 11.4288 | 0.1735 |
| **Fusiform** | 209.49 | 11.3824 | 0.1716 |
| **Isthmus Cingulate** | 207.72 | 11.5835 | 0.1680 |
| **Lateral Orbitofrontal** | 203.62 | 11.0561 | 0.1731 |
| **Superior Parietal** | 200.54 | 11.0222 | 0.1690 |
| **Superior Temporal Sulcus** | 200.27 | 11.3735 | 0.1696 |
| **Middle Temporal** | 199.48 | 11.2383 | 0.1714 |
| **Pars Triangularis** | 198.06 | 11.1687 | 0.1722 |
| **Rostral Middle Frontal** | 197.60 | 10.8487 | 0.1737 |
| **Inferior Temporal** | 196.40 | 11.1225 | 0.1723 |
| **Caudal Anterior Cingulate** | 196.28 | 11.4254 | 0.1700 |
| **Inferior Parietal** | 194.64 | 10.9589 | 0.1696 |
| **Superior Frontal** | 193.54 | 10.7116 | 0.1730 |
| **Lateral Occipital** | 193.06 | 11.1835 | 0.1694 |
| **Postcentral (S1)** | 191.64 | 11.1276 | 0.1703 |
| **Caudal Middle Frontal** | 190.88 | 10.9138 | 0.1726 |
| **Precuneus** | 188.53 | 11.1705 | 0.1672 |
| **Supra Marginal** | 187.49 | 11.2627 | 0.1700 |
| **Precentral** | 187.38 | 10.9731 | 0.1706 |
| **Pars Opercularis** | 182.56 | 11.1264 | 0.1709 |
| **Posterior Cingulate** | 181.68 | 11.1875 | 0.1688 |
| **Paracentral** | 180.86 | 10.7427 | 0.1697 |
| **Subcortex** | **Power** | | **Frequency** |
| | **% Increase** | **T Statistics** | **Hz** |
| **Hypothalamus** | 340.18 | 11.7229 | 0.1733 |
| **Basal Forebrain** | 268.41 | 12.0110 | 0.1732 |
| **Amygdala** | 265.19 | 11.5314 | 0.1728 |
| **Striatum** | 242.13 | 12.0794 | 0.1718 |
| **Medulla** | 239.55 | 11.0568 | 0.1715 |
| **Midbrain** | 239.41 | 11.8140 | 0.1717 |
| **Hippocampus** | 236.10 | 11.3878 | 0.1722 |
| **Thalamus** | 231.95 | 11.5374 | 0.1693 |
| **Pons** | 222.48 | 10.8429 | 0.1723 |
| **Cerebellum** | 221.25 | 10.7883 | 0.1735 |

**Table S3. Regional distributions of high-frequency BOLD oscillation.** BOLD power spectrogram was acquired on a voxel-level and the power spectrogram of each ROI was calculated as the normalized average across all voxels within this ROI. Based on the power spectrogram, we identified the spectral peaks in lower-frequency and higher-frequency ranges, traced the time courses of oscillation power, calculated the increase in oscillation power from wake to sleep, and the statistical difference (T value) in oscillation power between wake and sleep. Listed in the table are the spectral peak (Hz) and oscillation power (% increase from wake to sleep, statistical difference between wake and sleep) of high-frequency BOLD oscillation, averaged across two hemispheres and participants S1 to S36.

**Table S4. Number of voxels in individual ROIs**

| Fine atlas | |
|---|---|
| **Cortex** | **Number of Voxels** |
| **V1** | 487 |
| **MST** | 60 |
| **V6** | 86 |
| **V2** | 469 |
| **V3** | 346 |
| **V4** | 237 |
| **V8** | 70 |
| **4** | 399 |
| **3b** | 243 |
| **FEF** | 101 |
| **PEF** | 67 |
| **55b** | 91 |
| **V3A** | 116 |
| **RSC** | 87 |
| **POS2** | 157 |
| **V7** | 54 |
| **IPS1** | 89 |
| **FFC** | 146 |
| **V3B** | 49 |
| **LO1** | 51 |
| **LO2** | 52 |
| **PIT** | 74 |
| **MT** | 65 |
| **A1** | 58 |
| **PSL** | 109 |
| **SFL** | 129 |
| **PCV** | 114 |
| **STV** | 104 |
| **7Pm** | 63 |
| **7m** | 112 |
| **POS1** | 126 |
| **23d** | 79 |
| **v23ab** | 49 |
| **d23ab** | 59 |
| **31pv** | 64 |
| **5m** | 75 |
| **5mv** | 95 |
| **23c** | 127 |
| **5L** | 93 |
| **24dd** | 121 |
| **24dv** | 75 |
| **7AL** | 100 |
| **SCEF** | 150 |
| **6ma** | 162 |
| **7Am** | 138 |
| **7PL** | 70 |
| **7PC** | 111 |
| **LIPv** | 77 |
| **VIP** | 76 |
| **MIP** | 101 |
| **1** | 248 |
| **2** | 219 |
| **3a** | 118 |
| **6d** | 113 |

| | |
|---|---|
| **6mp** | 155 |
| **6v** | 104 |
| **p24pr** | 71 |
| **33pr** | 45 |
| **a24pr** | 62 |
| **p32pr** | 83 |
| **a24** | 84 |
| **d32** | 112 |
| **8BM** | 144 |
| **p32** | 69 |
| **10r** | 90 |
| **47m** | 60 |
| **8Av** | 202 |
| **8Ad** | 150 |
| **9m** | 256 |
| **8BL** | 152 |
| **9p** | 133 |
| **10d** | 157 |
| **8C** | 177 |
| **44** | 138 |
| **45** | 158 |
| **47l** | 130 |
| **a47r** | 216 |
| **6r** | 187 |
| **IFJa** | 80 |
| **IFJp** | 58 |
| **IFSp** | 104 |
| **IFSa** | 164 |
| **p9-46v** | 172 |
| **46** | 227 |
| **a9-46v** | 151 |
| **9-46d** | 240 |
| **9a** | 171 |
| **10v** | 135 |
| **a10p** | 109 |
| **10pp** | 111 |
| **11l** | 180 |
| **13l** | 106 |
| **OFC** | 149 |
| **47s** | 102 |
| **LIPd** | 50 |
| **6a** | 161 |
| **i6-8** | 98 |
| **s6-8** | 77 |
| **43** | 102 |
| **OP4** | 120 |
| **OP1** | 80 |
| **OP2-3** | 59 |
| **52** | 42 |
| **RI** | 73 |
| **PFcm** | 81 |
| **PoI2** | 114 |
| **TA2** | 78 |
| **FOP4** | 113 |
| **MI** | 97 |
| **Pir** | 70 |
| **AVI** | 92 |
| **AAIC** | 74 |

| | |
|---|---|
| **FOP1** | 53 |
| **FOP3** | 39 |
| **FOP2** | 41 |
| **PFt** | 106 |
| **AIP** | 124 |
| **EC** | 82 |
| **PreS** | 78 |
| **H** | 112 |
| **ProS** | 54 |
| **PeEc** | 182 |
| **STGa** | 84 |
| **PBelt** | 91 |
| **A5** | 135 |
| **PHA1** | 95 |
| **PHA3** | 80 |
| **STSda** | 124 |
| **STSdp** | 118 |
| **STSvp** | 136 |
| **TGd** | 334 |
| **TE1a** | 174 |
| **TE1p** | 229 |
| **TE2a** | 250 |
| **TF** | 191 |
| **TE2p** | 137 |
| **PHT** | 175 |
| **PH** | 157 |
| **TPOJ1** | 154 |
| **TPOJ2** | 129 |
| **TPOJ3** | 73 |
| **DVT** | 100 |
| **PGp** | 123 |
| **IP2** | 97 |
| **IP1** | 130 |
| **IP0** | 79 |
| **PFop** | 104 |
| **PF** | 219 |
| **PFm** | 328 |
| **PGi** | 285 |
| **PGs** | 206 |
| **V6A** | 45 |
| **VMV1** | 76 |
| **VMV3** | 57 |
| **PHA2** | 48 |
| **V4t** | 42 |
| **FST** | 86 |
| **V3CD** | 62 |
| **LO3** | 53 |
| **VMV2** | 43 |
| **31pd** | 57 |
| **31a** | 53 |
| **VVC** | 118 |
| **25** | 46 |
| **s32** | 41 |
| **pOFC** | 86 |
| **PoI1** | 72 |
| **Ig** | 47 |
| **FOP5** | 77 |
| **p10p** | 134 |

| | |
|---|---|
| **p47r** | 122 |
| **TGv** | 129 |
| **MBelt** | 65 |
| **LBelt** | 54 |
| **A4** | 119 |
| **STSva** | 92 |
| **TE1m** | 127 |
| **PI** | 61 |
| **a32pr** | 75 |
| **p24** | 72 |
| **Subcortex** | Number of Voxels |
| **Hippocampus** | 298 |
| **Amygdala** | 100 |
| **Basal Forebrain** | 167 |
| **Hypothalamus** | 46 |
| **Midbrain** | 391 |
| **Pons** | 860 |
| **Medulla** | 247 |
| **Thalamus (Anterior)** | 34 |
| **Thalamus (Lateral Dorsal)** | 51 |
| **Thalamus (Lateral Posterior)** | 26 |
| **Thalamus (Lateral Geniculate)** | 23 |
| **Thalamus (Pulvinar)** | 103 |
| **Thalamus (Parafascicular)** | 22 |
| **Thalamus (Centromedian)** | 29 |
| **Thalamus (Ventral Posterior)** | 40 |
| **Thalamus (Ventral Lateral)** | 79 |
| **Thalamus (Ventral Anterior)** | 29 |
| **Thalamus (Medial Dorsal)** | 51 |
| **Thalamus (Medial Geniculate)** | 26 |
| **Striatum (Limbic Network)** | 165 |
| **Striatum (Occipital Network)** | 35 |
| **Striatum (Parietal Network)** | 133 |
| **Striatum (Central Network)** | 103 |
| **Striatum (Temporal Network)** | 31 |
| **Striatum (Frontal Network)** | 386 |
| **Cerebellum (HIIV)** | 323 |
| **Cerebellum (HV)** | 378 |
| **Cerebellum (HVI)** | 817 |
| **Cerebellum (CrusI)** | 986 |
| **Cerebellum (CrusII)** | 662 |
| **Cerebellum (HVIIb)** | 260 |
| **Cerebellum (HVIIIa)** | 235 |
| **Cerebellum (HVIIIb)** | 140 |
| **Cerebellum (HIX)** | 221 |
| **Cerebellum (HX)** | 57 |
| **Cerebellum (Dentate)** | 158 |
| **Coarse atlas** | |
| **Cortex** | **Number of Voxels** |
| **Superior Temporal Sulcus** | 239 |
| **Caudal Anterior Cingulate** | 171 |
| **Caudal Middle Frontal** | 548 |
| **Cuneus (V2)** | 244 |
| **Entorhinal** | 114 |
| **Fusiform** | 701 |
| **Inferior Parietal** | 1144 |
| **Inferior Temporal** | 756 |
| **Isthmus Cingulate** | 217 |

| | |
|---|---|
| **Lateral Occipital** | 862 |
| **Lateral Orbitofrontal** | 629 |
| **Lingual (V2)** | 594 |
| **Medial Orbitofrontal** | 417 |
| **Middle Temporal** | 862 |
| **Parahippocampal** | 184 |
| **Paracentral** | 327 |
| **Pars Opercularis** | 420 |
| **Pars Orbitalis** | 207 |
| **Pars Triangularis** | 364 |
| **Pericalcarine (V1)** | 200 |
| **Postcentral (S1)** | 866 |
| **Posterior Cingulate** | 279 |
| **Precentral** | 1139 |
| **Precuneus** | 826 |
| **Rostral Anterior Cingulate** | 200 |
| **Rostral Middle Frontal** | 1402 |
| **Superior Frontal** | 1768 |
| **Superior Parietal** | 1099 |
| **Superior Temporal** | 920 |
| **Supra Marginal** | 853 |
| **Transverse Temporal (A1)** | 108 |
| **Insula** | 562 |
| **Subcortex** | Number of Voxels |
| **Hippocampus** | 298 |
| **Amygdala** | 100 |
| **Basal Forebrain** | 167 |
| **Hypothalamus** | 46 |
| **Midbrain** | 391 |
| **Pons** | 860 |
| **Medulla** | 247 |
| **Thalamus** | 499 |
| **Striatum** | 851 |
| **Cerebellum** | 3991 |

**Table S4. Number of voxels in individual ROIs.** We parcellated the brain into 42 coarse ROIs, including 32 cortical regions and 10 subcortical regions (cerebellum, striatum, thalamus, medulla, pons, midbrain, hypothalamus, basal forebrain, amygdala and hippocampus), or 217 fine ROIs, including 180 cortical regions, 11 cerebellum lobules, 7 striatum divisions, 12 thalamic subregions, medulla, pons, midbrain, hypothalamus, basal forebrain, amygdala and hippocampus. Listed in the table are the number of voxels in individual ROIs, summed over two hemispheres and averaged across participants S1 to S36.

**Table S5. Onset of BOLD oscillations**

| Onset of Low-frequency BOLD Oscillation | | | Onset of High-frequency BOLD Oscillation | | |
|---|---|---|---|---|---|
| | Lead Time | | | Lead Time | |
| | Mean (sec) | SEM (sec) | | Mean (sec) | SEM (sec) |
| **Thalamus (Lateral Geniculate)** | 12.517 | 0.621 | **Midbrain** | 9.405 | 0.550 |
| **Thalamus (Medial Geniculate)** | 11.833 | 0.708 | **Thalamus (Medial Geniculate)** | 7.907 | 0.905 |
| **Thalamus (Pulvinar)** | 8.884 | 0.345 | **Amygdala** | 7.631 | 0.338 |
| **Thalamus (Lateral Posterior)** | 8.533 | 0.492 | **Parahippocampal** | 7.197 | 0.307 |
| **Thalamus (Ventral Lateral)** | 8.250 | 0.466 | **Thalamus (Lateral Dorsal)** | 6.438 | 0.368 |
| **Lateral Occipital** | 8.116 | 0.410 | **Pons** | 6.194 | 0.445 |
| **Hippocampus** | 8.011 | 0.249 | **Thalamus (Anterior)** | 6.186 | 1.013 |
| **Cuneus (V2)** | 7.871 | 0.355 | **Thalamus (Medial Dorsal)** | 6.095 | 0.549 |
| **Lingual (V2)** | 7.850 | 0.297 | **Basal Forebrain** | 5.892 | 0.549 |
| **Precuneus** | 7.811 | 0.242 | **Cerebellum** | 5.711 | 0.286 |
| **Parahippocampal** | 7.787 | 0.329 | **Medulla** | 5.548 | 0.781 |
| **Pericalcarine (V1)** | 7.559 | 0.441 | **Isthmus Cingulate** | 5.516 | 0.297 |
| **Thalamus (Ventral Posterior)** | 7.171 | 0.420 | **Thalamus (Ventral Lateral)** | 5.217 | 0.402 |
| **Basal Forebrain** | 7.051 | 0.374 | **Pars Triangularis** | 5.167 | 0.298 |
| **Isthmus Cingulate** | 7.013 | 0.396 | **Hypothalamus** | 4.988 | 0.808 |
| **Superior Parietal** | 6.990 | 0.303 | **Transverse Temporal (A1)** | 4.904 | 0.384 |
| **Postcentral (S1)** | 6.963 | 0.195 | **Superior Frontal** | 4.771 | 0.303 |
| **Superior Temporal** | 6.421 | 0.227 | **Hippocampus** | 4.744 | 0.291 |
| **Inferior Parietal** | 6.403 | 0.350 | **Superior Temporal** | 4.543 | 0.203 |
| **Striatum** | 6.253 | 0.459 | **Medial Orbitofrontal** | 4.524 | 0.369 |
| **Superior Temporal Sulcus** | 6.212 | 0.390 | **Thalamus (Ventral Posterior)** | 4.443 | 0.349 |
| **Fusiform** | 6.050 | 0.299 | **Rostral Middle Frontal** | 4.296 | 0.354 |
| **Supra Marginal** | 5.862 | 0.273 | **Thalamus (Lateral Geniculate)** | 3.990 | 0.584 |
| **Posterior Cingulate** | 5.719 | 0.290 | **Rostral Anterior Cingulate** | 3.888 | 0.360 |
| **Thalamus (Ventral Anterior)** | 5.455 | 0.535 | **Precentral** | 3.635 | 0.204 |
| **Thalamus (Medial Dorsal)** | 5.451 | 0.457 | **Superior Parietal** | 3.583 | 0.294 |
| **Precentral** | 5.440 | 0.258 | **Lateral Occipital** | 3.463 | 0.254 |
| **Midbrain** | 5.411 | 0.254 | **Striatum** | 3.385 | 0.432 |
| **Thalamus (Anterior)** | 5.052 | 0.687 | **Pars Opercularis** | 3.240 | 0.209 |
| **Paracentral** | 4.880 | 0.347 | **Caudal Middle Frontal** | 3.180 | 0.237 |
| **Rostral Anterior Cingulate** | 4.819 | 0.345 | **Posterior Cingulate** | 3.101 | 0.296 |
| **Thalamus (Centromedian)** | 4.780 | 0.772 | **Lingual (V2)** | 3.032 | 0.246 |
| **Thalamus (Lateral Dorsal)** | 4.488 | 0.535 | **Cuneus (V2)** | 2.885 | 0.318 |
| **Thalamus (Parafascicular)** | 4.218 | 0.581 | **Postcentral (S1)** | 2.868 | 0.235 |
| **Pars Orbitalis** | 4.183 | 0.488 | **Lateral Orbitofrontal** | 2.847 | 0.181 |
| **Transverse Temporal (A1)** | 4.006 | 0.490 | **Fusiform** | 2.823 | 0.256 |
| **Insula** | 3.823 | 0.251 | **Inferior Parietal** | 2.812 | 0.259 |
| **Entorhinal** | 3.821 | 0.586 | **Thalamus (Pulvinar)** | 2.803 | 0.265 |
| **Lateral Orbitofrontal** | 3.727 | 0.396 | **Insula** | 2.611 | 0.251 |
| **Middle Temporal** | 3.647 | 0.200 | **Supra Marginal** | 2.509 | 0.281 |
| **Caudal Anterior Cingulate** | 3.356 | 0.391 | **Pericalcarine (V1)** | 2.341 | 0.314 |
| **Cerebellum** | 3.333 | 0.309 | **Thalamus (Parafascicular)** | 2.328 | 0.720 |
| **Pons** | 2.835 | 0.482 | **Paracentral** | 2.214 | 0.341 |
| **Caudal Middle Frontal** | 2.742 | 0.281 | **Inferior Temporal** | 2.122 | 0.260 |
| **Rostral Middle Frontal** | 2.679 | 0.347 | **Thalamus (Ventral Anterior)** | 2.101 | 0.664 |
| **Superior Frontal** | 2.678 | 0.390 | **Thalamus (Lateral Posterior)** | 2.095 | 0.655 |
| **Medial Orbitofrontal** | 2.499 | 0.401 | **Precuneus** | 1.979 | 0.271 |
| **Pars Triangularis** | 2.322 | 0.434 | **Pars Orbitalis** | 1.567 | 0.417 |
| **Inferior Temporal** | 2.252 | 0.310 | **Entorhinal** | 1.485 | 0.457 |
| **Amygdala** | 1.982 | 0.493 | **Middle Temporal** | 1.008 | 0.293 |
| **Medulla** | 1.146 | 0.608 | **Caudal Anterior Cingulate** | 0.781 | 0.279 |
| **Pars Opercularis** | 0.994 | 0.403 | **Superior Temporal Sulcus** | 0.105 | 0.470 |
| **Hypothalamus** | 0.000 | 0.717 | **Thalamus (Centromedian)** | 0.000 | 0.606 |

**Table S5. Onset of BOLD oscillations.** Cross-correlation analysis was applied to estimate the temporal lag between different brain regions in the onset of BOLD oscillations at the transition from wake to sleep. Listed in the table are the mean and standard error mean (SEM) of the lag structure averaged across two hemispheres and participants S1 to S36.

**Table S6. Offset of BOLD oscillations**

| Offset of low-frequency BOLD oscillation | | | Offset of high-frequency BOLD oscillation | | |
|---|---|---|---|---|---|
| | Lag Time | | | Lag Time | |
| | Mean (sec) | SEM (sec) | | Mean (sec) | SEM (sec) |
| Thalamus (Centromedian) | 0.000 | 0.622 | Thalamus (Parafascicular) | 0.000 | 0.530 |
| Thalamus (Anterior) | 1.622 | 0.472 | Hypothalamus | 0.255 | 0.472 |
| Thalamus (Parafascicular) | 2.014 | 0.423 | Thalamus (Lateral Posterior) | 0.826 | 0.493 |
| Medial Orbitofrontal | 2.221 | 0.224 | Thalamus (Ventral Posterior) | 1.318 | 0.437 |
| Pars Orbitalis | 2.814 | 0.271 | Thalamus (Lateral Geniculate) | 1.988 | 0.579 |
| Pars Triangularis | 2.867 | 0.178 | Thalamus (Medial Dorsal) | 2.168 | 0.418 |
| Thalamus (Medial Geniculate) | 3.111 | 0.367 | Thalamus (Lateral Dorsal) | 2.193 | 0.287 |
| Pars Opercularis | 3.218 | 0.231 | Amygdala | 2.222 | 0.405 |
| Rostral Middle Frontal | 3.361 | 0.492 | Pars Orbitalis | 2.424 | 0.338 |
| Caudal Middle Frontal | 3.493 | 0.457 | Rostral Middle Frontal | 2.542 | 0.283 |
| Hypothalamus | 3.752 | 0.657 | Striatum | 2.670 | 0.316 |
| Lateral Orbitofrontal | 3.919 | 0.247 | Basal Forebrain | 2.846 | 0.393 |
| Superior Parietal | 4.214 | 0.187 | Superior Frontal | 3.168 | 0.248 |
| Basal Forebrain | 4.266 | 0.415 | Medial Orbitofrontal | 3.286 | 0.298 |
| Lateral Occipital | 4.387 | 0.299 | Midbrain | 3.390 | 0.372 |
| Thalamus (Ventral Posterior) | 4.497 | 0.198 | Caudal Anterior Cingulate | 3.621 | 0.206 |
| Superior Temporal | 4.502 | 0.125 | Medulla | 3.707 | 0.382 |
| Hippocampus | 4.534 | 0.221 | Rostral Anterior Cingulate | 3.844 | 0.314 |
| Middle Temporal | 4.619 | 0.159 | Hippocampus | 3.977 | 0.315 |
| Superior Frontal | 4.630 | 0.482 | Lateral Orbitofrontal | 4.327 | 0.271 |
| Rostral Anterior Cingulate | 4.759 | 0.224 | Entorhinal | 4.583 | 0.197 |
| Fusiform | 4.871 | 0.144 | Pons | 4.617 | 0.258 |
| Midbrain | 4.882 | 0.325 | Thalamus (Pulvinar) | 4.826 | 0.247 |
| Pons | 4.976 | 0.355 | Posterior Cingulate | 5.265 | 0.204 |
| Precentral | 5.116 | 0.107 | Caudal Middle Frontal | 5.502 | 0.240 |
| Inferior Temporal | 5.206 | 0.185 | Transverse Temporal (A1) | 5.613 | 0.279 |
| Supra Marginal | 5.228 | 0.173 | Middle Temporal | 5.645 | 0.267 |
| Parahippocampal | 5.250 | 0.263 | Lateral Occipital | 5.820 | 0.239 |
| Thalamus (Pulvinar) | 5.358 | 0.171 | Thalamus (Centromedian) | 5.826 | 0.460 |
| Thalamus (Ventral Lateral) | 5.432 | 0.351 | Parahippocampal | 5.836 | 0.182 |
| Postcentral (S1) | 5.441 | 0.171 | Thalamus (Medial Geniculate) | 5.926 | 0.714 |
| Transverse Temporal (A1) | 5.522 | 0.211 | Superior Temporal | 5.980 | 0.191 |
| Thalamus (Medial Dorsal) | 5.574 | 0.364 | Superior Parietal | 5.998 | 0.236 |
| Entorhinal | 5.750 | 0.361 | Thalamus (Ventral Lateral) | 6.049 | 0.336 |
| Striatum | 5.828 | 0.288 | Cerebellum | 6.080 | 0.191 |
| Cerebellum | 5.857 | 0.406 | Insula | 6.145 | 0.223 |
| Amygdala | 5.894 | 0.428 | Postcentral (S1) | 6.153 | 0.208 |
| Insula | 5.983 | 0.160 | Pars Triangularis | 6.161 | 0.226 |
| Thalamus (Ventral Anterior) | 6.329 | 0.388 | Fusiform | 6.200 | 0.219 |
| Inferior Parietal | 6.357 | 0.318 | Thalamus (Anterior) | 6.228 | 0.678 |
| Thalamus (Lateral Geniculate) | 6.373 | 0.418 | Precentral | 6.229 | 0.219 |
| Caudal Anterior Cingulate | 6.419 | 0.264 | Inferior Temporal | 6.296 | 0.232 |
| Thalamus (Lateral Dorsal) | 6.487 | 0.294 | Lingual (V2) | 6.403 | 0.221 |
| Lingual (V2) | 6.520 | 0.306 | Pars Opercularis | 6.523 | 0.236 |
| Medulla | 6.681 | 0.326 | Isthmus Cingulate | 6.598 | 0.190 |
| Pericalcarine (V1) | 6.724 | 0.380 | Supra Marginal | 6.637 | 0.241 |
| Superior Temporal Sulcus | 7.045 | 0.245 | Inferior Parietal | 6.707 | 0.236 |
| Posterior Cingulate | 7.439 | 0.283 | Superior Temporal Sulcus | 6.856 | 0.252 |
| Isthmus Cingulate | 7.529 | 0.285 | Paracentral | 6.938 | 0.273 |
| Paracentral | 7.541 | 0.341 | Precuneus | 7.044 | 0.255 |
| Precuneus | 7.648 | 0.314 | Cuneus (V2) | 7.426 | 0.326 |
| Cuneus (V2) | 7.744 | 0.521 | Thalamus (Ventral Anterior) | 7.540 | 0.611 |
| Thalamus (Lateral Posterior) | 10.030 | 0.377 | Pericalcarine (V1) | 7.821 | 0.284 |

**Table S6. Offset of BOLD oscillations.** Cross-correlation analysis was applied to estimate the temporal lag between different brain regions in the offset of BOLD oscillations at the transition from sleep to wake. Listed in the table are the mean and standard error mean (SEM) of the lag structure averaged cross two hemispheres and participants S1 to S36.
